# Data- and knowledge-derived functional landscape of human solute carriers

**DOI:** 10.1101/2024.10.14.618155

**Authors:** Ulrich Goldmann, Tabea Wiedmer, Andrea Garofoli, Vitaly Sedlyarov, Manuel Bichler, Gernot Wolf, Eirini Christodoulaki, Alvaro Ingles-Prieto, Evandro Ferrada, Fabian Frommelt, Shao Thing Teoh, Philipp Leippe, Ben Haladik, Gabriel Onea, Martin Pfeifer, Mariah Kohlbrenner, Lena Chang, Paul Selzer, Jürgen Reinhardt, Daniela Digles, Gerhard F. Ecker, Tanja Osthushenrich, Aidan MacNamara, Anders Malarstig, David Hepworth, Giulio Superti-Furga

## Abstract

Research on the understudied solute carrier (SLC) superfamily of membrane transporters would greatly profit from a comprehensive knowledgebase, synthesizing data and knowledge on different aspects of SLC function. We consolidated multi-omics data sets with selected curated information from the public domain, such as structure prediction, substrate annotation, disease association and subcellular localization. This SLC-centric knowledge is made accessible to the scientific community via a web portal, featuring interactive dashboards and a tool for family-wide, tree-based visualization of SLC properties. Making use of the systematically collected and curated data sets, we selected eight feature-dimensions to compute an integrated functional landscape of human SLCs. This landscape represents various functional aspects, harmonizing local and global features of the underlying data sets, as demonstrated by inspecting structural folds and subcellular locations of selected transporters. Based on all available data sets and their integration, we assigned a biochemical/biological function to each SLC, making it one of the largest systematic annotations of human gene function and likely acting as a blueprint for future endeavours.

## Introduction

Systematically assessing human gene function is an important but daunting challenge in any defined biological system. The RESOLUTE and REsolution consortia mounted a concerted action to tackle the solute carrier (SLC) superfamily of membrane transporters, a subsystem thought to be approachable by size (∼ 450 genes), of coherent biochemical role and related as well as interconnected functions, spanning the entire functional space of secondary active transport of small organic and inorganic molecules. SLCs have a special role in pharmacology as they include targets of some very prominent drugs, like gliflozins and monoamine uptake inhibitors (Wang *et al*, 2020; Galetin *et al*, 2024), contain many disease-linked genes (Lin *et al*, 2015; Wang *et al*, 2020; Schlessinger *et al*, 2023), and fulfil important functions in drug uptake and disposition (Girardi *et al*, 2020; Giacomini *et al*, 2010). It is not surprising, therefore, that interest around SLC transporters has increased in the pharmaceutical industry (César-Razquin *et al*, 2015). The RESOLUTE consortium of the Innovative Medicine Initiative of the European Union was a public-private partnership that started in 2018 involving seven pharmaceutical companies and six academic institutions aimed at broadly unlocking solute carriers as a target class. The goal was to provide research tools and advance the knowledge for as many human SLCs as possible, thus lowering the entry barrier for further research projects and drug discovery ((Superti-Furga *et al*, 2020); see also https://re-solute.eu/communications for a detailed account of the project). In 2021, the REsolution consortium formed with nine partners to extend the scope of RESOLUTE by anchoring its results in a biomedical context (Wiedmer *et al*, 2022). The challenges were manifold. How to tackle the experimental functional annotation of a group of genes of such size and scope? How to optimize knowledge advancement in the most efficient way? How to integrate data to achieve knowledge beyond the sum of the parts?

In the RESOLUTE consortium we focused on as few experimental systems as possible, and employed robust, parallelizable and multiplexable approaches to synergize the advancement of knowledge. Solute carriers are a superfamily of transporters, arbitrarily grouped to rationalize the genetic nomenclature for multipass transmembrane carriers that are not ATPases, aquaporins or channels (Hediger *et al*, 2004). SLCs localize to the plasma membrane and different intracellular membranes (Pizzagalli *et al*, 2021). They have been classified into 70 families, with family members sharing more than 20% sequence identity (Hediger *et al*, 2004). Recently, AlphaFold enabled to discern 24 different structural folds to which the families can be allocated (Ferrada & Superti-Furga, 2022). Many of these folds are evolutionary ancient, reflecting their fundamental cellular and physiological role (Höglund *et al*, 2011). As a result of different rounds of gene duplication, paralog genes have contributed to extensive functional redundancy. This is part of the reason why assigning specific functions to single SLC genes has not been straightforward and a good proportion of SLC gene could be termed ‘orphan’ (César-Razquin *et al*, 2015; Meixner *et al*, 2020). SLCs are expressed across all human tissues but their specificity of expression varies widely: while some SLCs are expressed only in very specific tissues, others are ubiquitously expressed across all tissues (César-Razquin *et al*, 2015; Zhang *et al*, 2019; O’Hagan *et al*, 2018). Each cell and tissue expresses around half of all SLCs, which is thought to reflect the metabolic requirements of that particular tissue (César-Razquin *et al*, 2015; O’Hagan *et al*, 2018). Of these, only between 20-30 SLCs are deemed essential (Wang *et al*, 2015; Blomen *et al*, 2015). The 13 glucose transporters within the SLC2 and SLC5 families provide an exemplary case of how the systemic redundancy, discrepancy in essentiality and sheer number of SLC genes challenge traditional cell-biological and biochemical approaches to gene function assignments.

It can be expected that most small molecules transported by the ∼ 450 members of the SLC superfamily are endogenous metabolites, from inorganic ions to complex prosthetic groups. However, some transporters may have evolved to transport environmental components that have been endogenously utilized in the past but turned into xenobiotics (Gründemann *et al*, 2005). Nevertheless, the chemical space for which transport has evolved is finite, with the human metabolome database currently listing ∼220,000 chemical species (https://hmdb.ca). The number of genes encoding proteins responsible for transport of molecules, other than water or the ions associated with channels, is relatively small, estimated to be under a thousand (Elbourne *et al*, 2017). SLCs represent the largest group among these. From these considerations, the problem of assigning functions to each of the SLC genes is one of matchmaking. Such a SLC-solute matchmaking process can be facilitated by exclusion, systematically eliminating unlikely pairings based on experimental data, known gene characteristics (also from other organisms), and functional or structural constraints. By ruling out non-functional or less likely associations, the set of possible gene-function matches is progressively narrowed down, ultimately leading to an optimal mapping where each gene is matched to its most likely or confirmed function. Such an approach relies on both the inclusion of positive matches and the exclusion of incompatible or improbable ones, ensuring a precise and accurate assignment of functions to genes.

Thus, the effort of creating a functional landscape is, essentially, optimising a virtual functional distance between SLC genes using data modalities we obtained in the RESOLUTE and REsolution efforts. Among the experimental data modalities targeted metabolomics provides the most direct assessment of substrate transport and its implications on metabolic consequences (Wiedmer *et al*, 2024). It is complemented with transcriptomics to the cellular functional state (Wiedmer *et al*, 2024) and with interaction proteomics providing the protein environment as molecular basis for functional states (Frommelt *et al*, 2024). Knowledge of the subcellular localization of SLCs further complements the understanding of their cellular environment. In this study we obtained experimental evidence of subcellular localization for the entire SLC superfamily. Beyond experimental data, we collected publicly available knowledge of both physical and physiological parameters relevant for and contributing to SLC function and further curated them for integration: structure prediction, tissue expression as well as substrate annotation and disease association.

Integration of biomedical, multimodal data sets remains challenging (Cai *et al*, 2022). A number of methods have been developed, each with specific requirements and applications. Multi-omics factor analysis (MOFA) (Argelaguet *et al*, 2018) and MultiMAP (Jain *et al*, 2021) are tailored for multi-omics data. They do not require complete data, but they will not work with qualitative data such as disease associations or lists of annotated substrates. Translating these annotations to quantitative similarities allows application of different integration algorithms such as similarity network fusion (SNF) (Wang *et al*, 2014) or AlignedUMAP (Dadu *et al*, 2023). They do, however, require complete data and will not work with missing data. In the present study we propose and apply a new approach to integrate the diverse data sets collected to a functional landscape of human SLCs.

It should be noted that there is no single right solution as there are many more transported molecules than transporters and the degree of redundancy in the SLC-solute matchmaking increases with our knowledge on them. Accordingly, any concept of ‘deorphanisation’, familiarly used in reference to nuclear hormone receptors and GPCRs, would be misleading in the case of SLC transporters. How then can a function of an SLC gene be assigned in an unequivocal way? The present study represents such an effort.

## Results

### Characterization of the human SLC superfamily members by their structure, expression and substrates

To start a systematic assignment of biochemical and biological properties to each member of the solute carrier superfamily demanded a definition of the criteria used to include or exclude genes from the analysis. To date, the HUGO Gene Nomenclature Committee (HGNC) included 449 human genes in the solute carrier superfamily (Seal *et al*, 2023). An additional fourteen genes were suggested by Perland & Fredriksson and another one by Ferrada & Superti-Furga (TMEM144) (Perland & Fredriksson, 2017; Ferrada & Superti-Furga, 2022), making up a total of 464 human genes (**Supplemental Table 1**). These included eight pseudogenes, two insertases (MTCH1 and MTCH2) and three auxiliary subunits (SLC3A1, SLC3A2, SLC51B) without transport function. Based on the sequence similarity of SLCs, the superfamily was divided into 70 families (Perland & Fredriksson, 2017; Hediger *et al*, 2004). At the start of the RESOLUTE consortium, a slightly lower number of proteins were classified as SLCs and therefore only 447 SLCs of 69 families were considered for the collection of data in this study.

Based on experimental structures and structures modelled using AlphaFold, we have recently identified 24 distinct transmembrane structural folds within the SLC superfamily (Ferrada & Superti-Furga, 2022). Inspired by the description and illustration of the human kinome (Manning *et al*, 2002), a similarly large and important protein superfamily, we constructed an unrooted tree, clustering the SLCs based on their structural similarity. The SLC tree visualized the folds in different colours (**Fig. 1A**), and a web-based version allowed interactive navigation down to individual SLCs as well as the upload and overlay with any gene-level annotation (https://re-solute.eu/resources/dashboards/slctree). We considered this tree as consisting of five major branches. Four of these were the MFS, LeuT, MitC and DMT clades, each with more than 30 members of a common structural fold and collectively covering two thirds of all SLCs (n=301; 67.3%). The fifth branch, covering one third of the SLCs (n=146; 32.7%), was much more heterogeneous and represented 20 more structural folds with less than 25 members each and also quite different numbers of transmembrane helices (UraA, Glt, NhaA, ZIP, SLC64, IT, MnN3, SLC51, SLC53, SLC44, SLC56, AmtB, SLC41, PiT, MATE, NPC1, CNT2, CNT1, YiiP, NCX). Like the kinome tree, the SLC tree offered itself as visualisation platform for a variety of SLC properties and data useful to be annotated from an evolutionary structural perspective, such as substrate preference.

**Figure 1.**
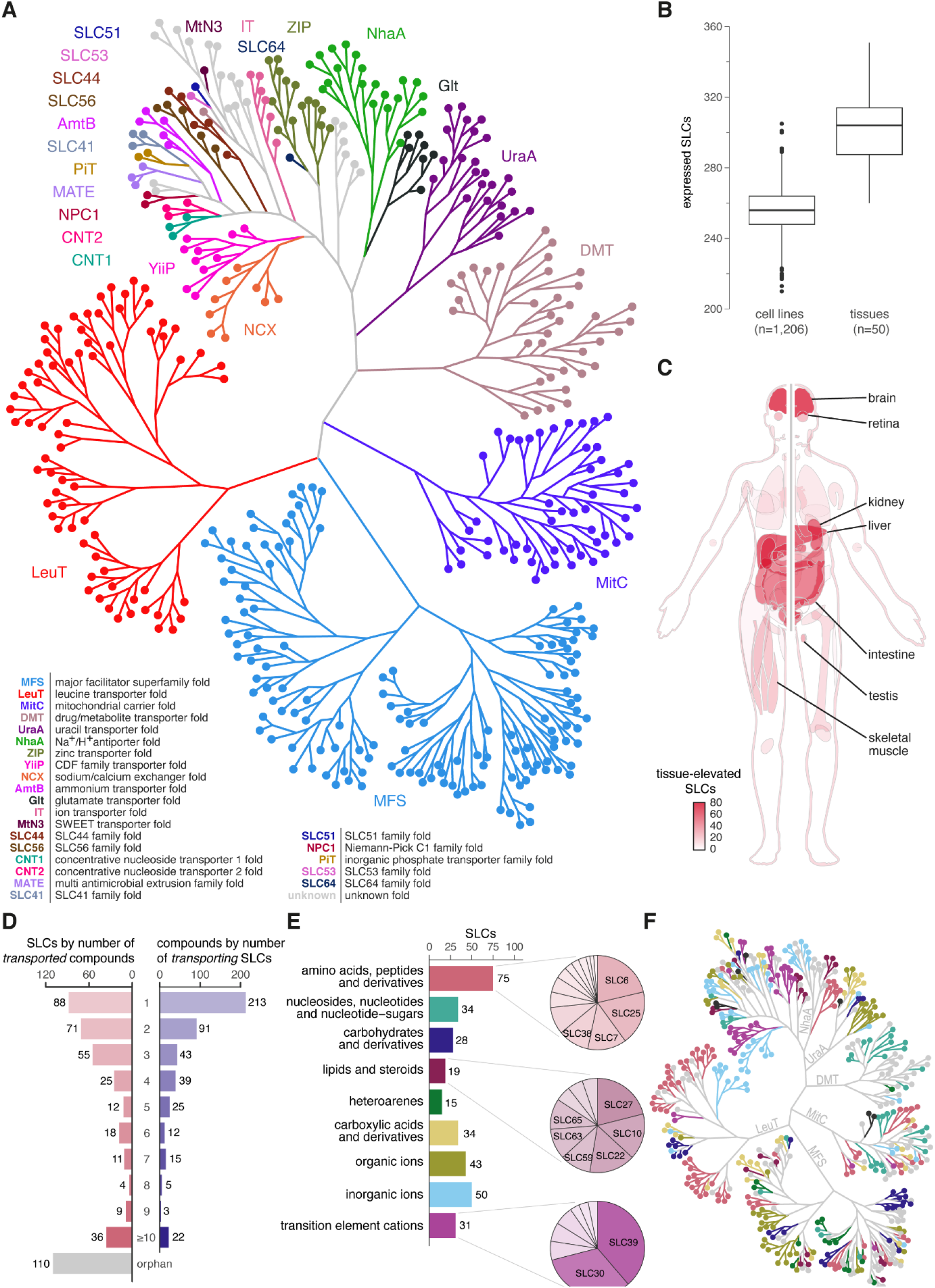
Overview of the solute carrier superfamily. **(A)** Unrooted tree representation of the 447 SLCs included in this study, based on experimental and modelled structures (data from (Ferrada & Superti-Furga, 2022)). The SLC tree illustrates 24 folds based on their similarity in different colours. A web-based version allows its exploration down to individual SLCs as well as uploading and overlaying with gene-level annotations (https://re-solute.eu/resources/dashboards/slctree). **(B)** Distribution of the numbers of expressed SLC genes within 1,206 cell lines and 50 tissues (data from (Uhlén *et al*, 2015)). **EV** Fig. 1A and **EV** Fig. 1B show the number of expressed SLC genes also in relation to the total number of expressed genes in a particular cell line or tissue, respectively. **(C)** Schematic representation of a human body where tissues are coloured by number of SLCs with a corresponding tissue-elevated expression pattern. Tissues featuring more than 20 SLCs are indicated. **EV** Fig. 1C shows the data for all tissues. **(D)** Summary of literature-based substrate annotation, demonstrating the redundancy of compounds transported per SLC as well as the redundancy of SLCs transporting a specific compound. The left chart shows the numbers of SLCs annotated to transport from 1 to 10 or more substrates, and SLCs without annotated substrate (i.e. orphan). The right chart shows the number of compounds annotated to be transported by 1 to 10 or more SLCs. **(E)** Bar chart with the number of SLCs classified into nine substrate classes. Pie charts illustrate the proportion of families within SLCs annotated as ‘amino acids, peptides and derivatives’, ‘lipids and steroids’ and ‘transition element cations’ transporters. **(F)** Substrate class for each SLC illustrated on the SLC tree. Colour code according to substrate class as in panel E.

The SLC family did not only involve a variety of structural folds, but also displayed various patterns of expression. Human Protein Atlas provided cell line and tissue expression data for 455 human SLCs (**Fig. 1B**) (Uhlén *et al*, 2015). Within the 1,206 cell lines analysed, we found an average of 2.2% of the genes expressed being SLCs. An individual cell line expressed 256 SLCs (56.2%) on average (**EV Fig. 1A**). The 50 tissues analysed also featured an average of 2.2% of the genes expressed being SLCs. Tissues had an average of 300 SLCs (65.9%) expressed (**EV Fig. 1B**). Human Protein Atlas also defined ‘tissue-elevated’ gene expression as a gene having elevated expression levels in a single tissue or a group of tissues (**EV Fig. 1C**). The group of brain tissues showed the highest number of tissue-elevated SLCs (n=72), followed by liver (n=62), kidney (n=53), the group of intestine tissues (n=47), testis (n=31), retina (n=25) and skeletal muscle (n=21) (**Fig. 1C**).

The complex structural categorization and the diverse expression patterns within this superfamily of proteins are likely rooted in their ability to transport a wide array of substances across different cellular membranes. We previously generated a comprehensive overview of substrates of the entire superfamily through manual annotation based on literature (Meixner *et al*, 2020). Considering recent advances of the field and limitations of the previous annotation, we updated and refined the annotation, as well as the inclusion criteria and the mapping process (Methods). This resulted in 2,044 SLC-compound-publication pairs. Every SLC-substrate annotation was matched to a ChEBI term (Hastings *et al*, 2016) and had at least one reference to a primary resource, mainly via PubMed identifier. The annotation is based on 678 publications, assigning 468 distinct substrates to 329 SLCs, with a median of 3 substrates transported per SLC. About a quarter of these SLCs (n=88; 26.7%) had only a single substrate annotated while 36 SLCs (10.9%) were annotated with more than ten annotated substrates each, of which SLC22A1 had the most annotations with 39 substrates, illustrating the heterogeneity in both substrate specificity but also knowledge across this transporter family. A quarter of the whole family (110 SLCs) remained without an annotated substrate (i.e. orphans) (**Fig. 1D**). Compared to the previously published annotation (Meixner *et al*, 2020), 23 SLCs now have substrates assigned and another 13 SLCs with previously annotated substrates were now classified as orphan since existing evidence did not fulfil our set criteria (**EV Fig. 1D**). The 468 distinct compounds described in this annotation were transported by a median of 2 SLCs. While nearly half of the compounds (n=213; 45.5%) were transported by only a single SLC, another third (n=164, 35.0%) of the compounds had three or more known transporters. In the future, such information may be used in rationalizing the choice of drug targets, acknowledging that some SLCs may have more diffuse effects due to their relative promiscuity.

To enable SLC superfamily-wide analyses of SLC substrate specificity and to facilitate integration with other data sets, we established a single-label classification of all substrates for each SLC. We defined a set of nine biologically and biochemically relevant substrate classes based on a manual selection of higher-level ChEBI terms (‘amino acids, peptides and derivates’; ‘carbohydrates and derivatives’; ‘carboxylic acids and derivatives’; ‘heteroarenes’; ‘inorganic ions’; ‘lipids and steroids’; ‘nucleosides, nucleotides and nucleotide-sugars’; ‘organic ions’; ‘transition element cations’). Using the ChEBI ontology tree, the substrates were matched to these substrate classes. Substrates that matched different classes were semi-automatically assigned to a single class. Similarly, SLCs with multiple substrates of different classes were assigned to a single class. Orphan SLCs remained the largest group with 110 SLCs. Among the 329 substrate-annotated SLCs, ‘amino acids, peptides and derivates’ was the largest substrate class (75 SLCs; 22.8%), followed by ‘inorganic ions’ (50 SLCs; 15.2%) and ‘organic ions’ (43 SLCs; 13.1%) (**Fig. 1E**). Most structural folds featured SLCs of diverse substrate classes (**Fig. 1F**).

Using this ontology-based approach to classify substrates, we were able to divide the largest group ‘other’ of unclassified substrates from the previous substrate classification (Meixner *et al*, 2020) into biologically and biochemically relevant substrate classes (**EV Fig. 1D**). In addition, SLCs from the previous groups ‘ion’ and ‘metal’ were split into more precisely termed groups, mostly into ‘carboxylic acids and derivatives’, ‘organic ions’, ‘inorganic ions’ and ‘transition element cations’.

### Survey of SLC superfamily-wide disease associations

The diversity of the substrates of the entire SLC superfamily and their resulting linkage to broad metabolic processes, implies a broad role in physiology in general and human health in particular. Many SLCs are linked to Mendelian diseases, even more are associated to a large variety of human disease traits, and several members were shown to be successful drug targets (Lin *et al*, 2015; Wang *et al*, 2020; Schlessinger *et al*, 2023). To obtain a comprehensive overview of the current knowledge of SLC genetics and its impact on human biology, we performed programmatic data mining to collect and curate data from a wide range of open-source databases (**Fig. 2A**). Selected databases included genetic and clinical evidence (UniProt, Ensembl, gnomAD, ClinVar, Orphanet) and statistically processed data (IEU OpenGWAS project, Genebass, Open Targets, Priority index) (**Supplemental Table 2**).

**Figure 2.**
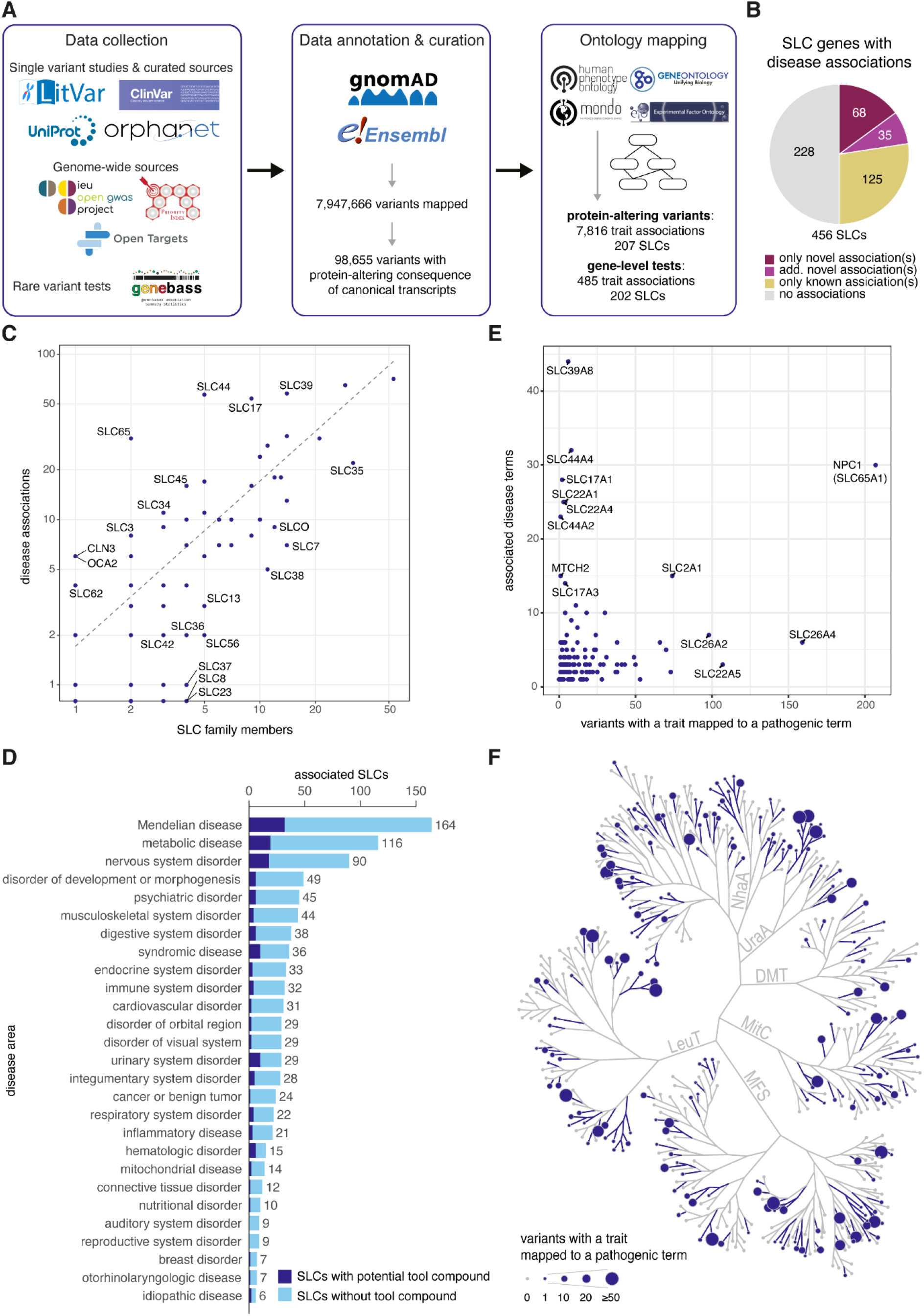
Collection of SLC genetic variants and disease associations. **(A)** Overview of workflow, consisting of data collection, data annotation & curation and ontology mapping. **(B)** Proportion of SLC genes with disease associations when comparing current data set to previous pathogenic associations in ClinVar and Orphanet. ‘Only novel associations’: no previous pathogenic associations in respective sources; ‘additional novel associations’: novel associations in addition to previously reported pathogenic associations in respective sources; ‘only known associations’: our data set includes the same pathogenic associations as previously reported by respective sources. **(C)** Pathogenic associations per SLC family. Highlighted families have more than twice or less than half the associations expected based on the average 1.7 pathogenic trait associations per SLC gene (dashed line). **(D)** Distribution of pathogenic associations of SLCs across main disease areas (≥ 5 associated SLCs). Disease areas are defined by the child terms of the Mondo term ‘Human disease’ (Methods). Only the disease areas ‘hematologic disorder’ and ‘urinary system disorder’ have more than one third of their associated SLCs covered by potential tool compounds, as defined in (Digles *et al*, 2024). **(E)** Number of distinct associated diseases compared to overall number of pathogenic variants in each SLC. **(F)** The number of pathogenic variants per SLC highlighted on the SLC tree.

Our methodical approach resulted in a curated data set consisting of 456 members of the SLC superfamily, offering a unified repository of information previously dispersed across various resources. We found 7,947,666 variants mapped to human SLC genes as well as various genetic and phenotypic annotations. These variants are in large proportion neutral but can also be deleterious through various types of mechanisms including direct impact on gene expression, protein structure or function. A protein-altering consequence on the canonical transcript of an SLC gene was annotated for 98,655 of those variants. Missense variants, which are single nucleotide mutations translating into single amino acid substitutions, made up 90.1% of all the collected protein-altering variants (**EV Fig. 2A**).

The compiled data set encompassed a wide range of associated traits, including human diseases, symptoms or biological quantitative measurements and general phenotypes not directly linked to pathogenicity. Associated traits were curated to a standardized terminology to eliminate redundancies, clarify ambiguous terms and distinguish disease-related from non-disease traits. We implemented a workflow to map terms to community-established ontologies, employing Monarch Disease Ontology (Mondo) (Vasilevsky *et al*, 2022) for disease association and the Experimental Factor Ontology, Human Phenotype Ontology, or Gene Ontology for broader biological terms (Malone *et al*, 2010; Köhler *et al*, 2021; Ashburner *et al*, 2000). The process of standardization and ontology mapping enabled us to annotate previously unclassified variants as likely pathogenic due to their association to disease traits.

Among the protein-altering variants, we found across 207 SLCs a total of 7,816 associations with traits, of which 3,244 (41.5%) were classified pathogenic. Moreover, 485 trait associations for 202 SLCs emerged from bundled missense and loss-of-function (LoF) gene-level tests for rare variants, of which 216 (44.5%) were classified pathogenic. Overall, we collected and mapped 780 distinct associations of pathogenic terms to 228 SLCs (**Fig. 2A, EV Fig. 2B**). For the following analyses we focused on this set of 3,244 pathogenic associations of protein-altering variants in canonical SLC transcripts (variant-level) or to the set of 780 merged pathogenic gene-level associations (gene-level).

By distinguishing between pathogenic variant-associations that are clinically established (defined as the ones reported in the ClinVar or Orphanet data sets), and the associations collected in our study that also contain likely pathogenic ones, we found that our data set contained only the reported associations for 125 SLCs. Beyond those, we found new likely pathogenic associations for 103 SLCs, with 68 of them without any previous association in ClinVar or Orphanet (**Fig. 2B**). An overview of those 68 SLCs and their newly associated 127 likely pathogenic terms revealed for few SLCs associations to multiple different pathogenic terms (e.g. SLC17A1, SLC22A1, SLC44A2, MTCH2) but for most of them less than five associations (**EV Fig. 2C**). Among them we found the association of systemic lupus erythematosus (SLE) with SLC15A2, SLC15A4 and SLC17A3. Although the involvement of SLC15A4 in the development of SLE has been well-described (Kobayashi *et al*, 2014; Heinz *et al*, 2020), these missense variants were not yet classified as pathogenic in ClinVar and Orphanet. No reports exist to date for the pathogenicity of SLC15A2 and SLC17A3 in SLE. Furthermore, type II diabetes was associated with SLC16A11, SLC30A8, SLC39A11 and MTCH2. The associations of SLC30A8 and SLC16A11 with diabetes are well-described but the disease mechanisms have not been fully elucidated yet (Krentz & Gloyn, 2020; Rusu *et al*, 2017; Hoch *et al*, 2019) and accordingly ClinVar and Orphanet do not report this association. MTCH2 was among the genes with many novel likely pathogenic associations, which have been previously reported in genome-wide association studies focused on obesity and diabetes (Kang *et al*, 2020). In contrast, we found no literature about the role of SLC39A11 in type II diabetes.

The new associations uncovered by our analysis suggest that other SLCs with already known pathogenic protein-altering genetic alterations, might still have incomplete annotations. Overall, we found an average of 1.7 pathogenic trait associations per SLC gene. Families SLC44, SLC17 and SLC39 showed exceptionally high numbers of pathogenic associations, each with more than 90 associations (**Fig. 2C**).

To get an overview of the implications of the superfamily’s disease associations, we mapped them to different disease areas employing the Mondo ontology (Methods). The disease areas with most SLCs associated were ‘Mendelian disease’ (164 SLCs), ‘metabolic disease’ (116 SLCs) and ‘nervous system disorder’ (90 SLCs) (**Fig. 2D**). Considering the list of potential tool compounds published recently for 51 SLCs (Digles *et al*, 2024) revealed that only few disease areas are somewhat covered with potential tool compounds for their associated SLCs. An SLC was associated with four disease areas on average, and only few SLCs associated with considerably more diseases, such as: SLC39A8, SLC44A4, NPC1, SLC17A1, SLC22A1, SLC22A4, and SLC44A2, each with more than 20 associated disease terms (**Fig. 2E**). SLC39A8 is a well-documented case of a pleiotropic gene (i.e. one gene associated with multiple phenotypes), for which only six pathogenic protein-altering variants have been linked to 44 different disease terms in humans (Pickrell *et al*, 2016). Overall, our analysis of human genetic data identified 780 disease associations for 228 SLCs. At the variant level, this represents 2,402 pathogenic variants (**Fig. 2F**) and 138 previously unknown likely pathogenic alleles.

### Experimental annotation of subcellular location for the SLC superfamily

The subcellular localization of SLCs is a functional property that is to date rather broadly annotated in public databases. But those data sets have limitations and caveats, such as for example annotation based on sequence similarity or antibody-based experimental evidence even though antibodies often lack a thorough validation of their target specificity (Ayoubi *et al*, 2023). Therefore, we collected and generated and curated experimental evidence for each member of the SLC superfamily.

We processed 560 Jump-In T-REx HEK293 cell lines, each expressing an SLC fused to an HA-tag under a doxycycline-inducible promoter followed by an immunofluorescence-based high-content imaging approach (**Fig. 3A**). Of those, 447 were cell lines expressing SLCs HA-tagged at the C-terminus and 113 at the N-terminus, the latter due to unsatisfactory C-terminal tagging. SLC expression was induced by doxycycline, followed by a washout period and subsequent immunofluorescent staining with HA and compartment markers for cytoplasm, plasma membrane, mitochondria, lysosome, endosome, endoplasmic reticulum (ER), Golgi, peroxisome, cytoskeleton and the nucleus. Using high-content imaging we acquired at least 768 images per cell line, that were SLC- and compartment-wise individually levelled and then visually inspected. We generated a manual assessment of ten consolidated locations (ER; Golgi; plasma membrane; perinucleic region; lysosomes; endosomes; mitochondria; cytosol; cytoskeleton; lamellipodia) for 407 SLCs that were adequately expressed. Each assessment contained for each compartment (i) a signal score representing the approximated relative fluorescence intensity compared to the total fluorescence intensity, and (ii) a confidence value ranging from 1 (very weak) to 5 (very high). The annotations were carefully curated in several rounds to minimize human bias and provide a ground truth for the development of machine learning based localization prediction. The last curation rounds were focussing on studying mismatches between prediction and human assessment leading to both, an improved human statement as well as improved predictive models (Baranowski *et al*, in preparation).

**Figure 3.**
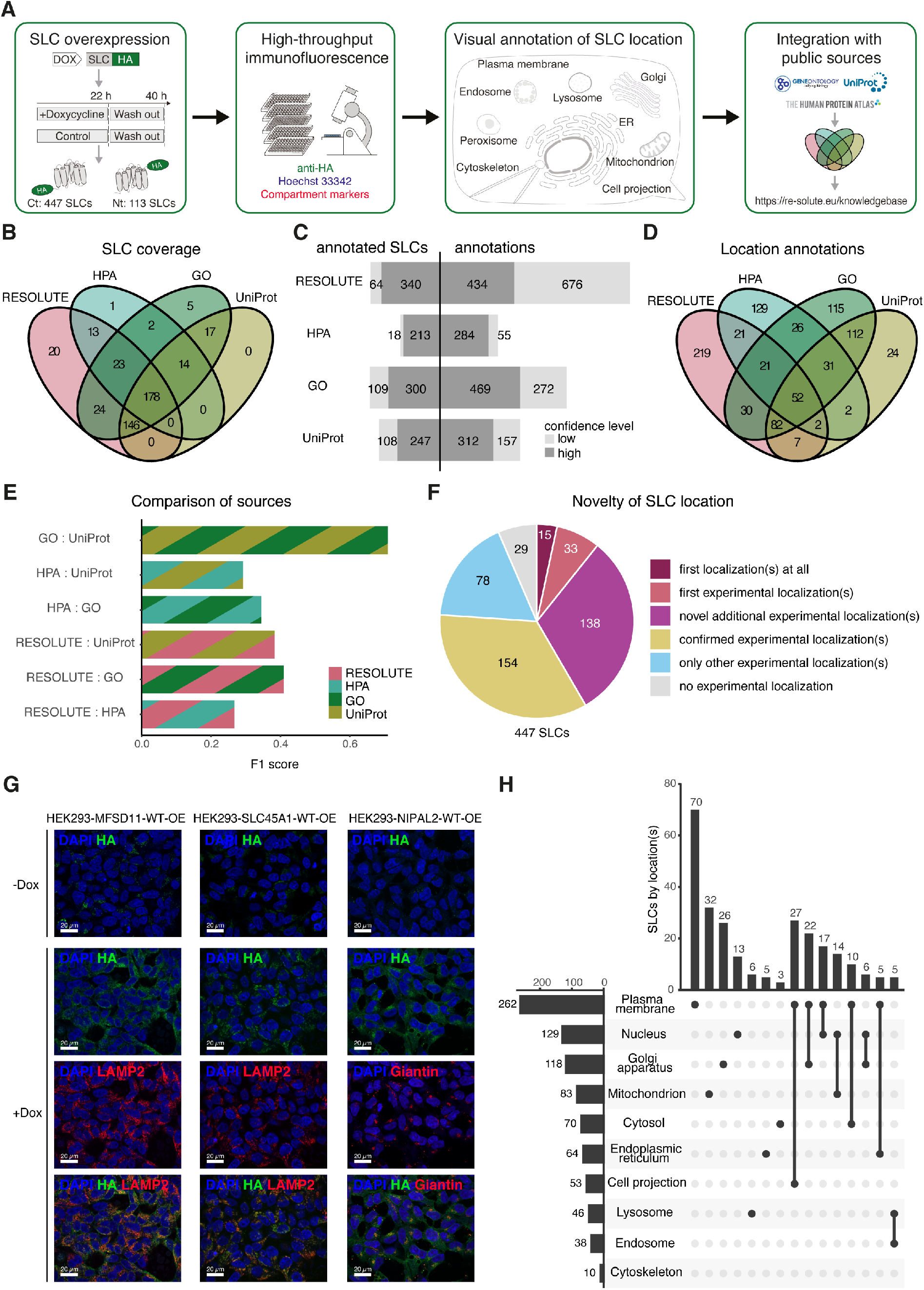
Subcellular localization of the SLC superfamily. **(A)** Workflow for the generation of a subcellular localization data set by experimental high-throughput immunofluorescence combined with integration of public data. **(B)** Comparison of SLC coverage between the acquired data set (RESOLUTE) and public sources Human Protein Atlas (HPA), Gene Ontology (GO) and UniProt. **(C)** Numbers of total and high-confidence annotated SLCs and localization annotations by RESOLUTE, HPA, GO and UniProt. For filtering criteria please refer to the Methods section. **(D)** Comparison of the overlap between high-confidence localization annotations in RESOLUTE, HPA, GO and UniProt. **(E)** Scoring of the overall agreement in high-confidence subcellular localization annotations for each pair of annotation sources. The F1-score combines precision and recall for each localization, with a higher score corresponding to higher consistency. **(F)** Novelty of high-confidence localization annotation per SLC by the RESOLUTE data set compared to HPA, GO and UniProt. **(G)** Immunofluorescence images of cell lines HEK293- MFSD11-WT-OE, HEK293-SLC45A1-WT-OE and HEK293-NIPAL2-WT-OE. Cells were treated with doxycycline for 22h followed by an 18h wash out. Green channel: HA-tag of overexpressed SLC. Red channel: compartment marker for lysosomes (LAMP2) or Golgi (Giantin). Blue channel: Hoechst 33342 for nucleus. Images were acquired using a 60x objective. Scale bars indicate 20 µm. **(H)** UpSet plot (Conway *et al*, 2017) visualizing the number of SLCs annotated for each location (horizontal bars) and number of SLCs annotated for specific location combinations (vertical bars) in the consolidated high-confidence data set with combined evidence from RESOLUTE, HPA, GO and UniProt. Location combinations with less than 5 SLCs or more than two locations are not shown.

To compare and complement our experimental results and annotations of subcellular localization with publicly available sources, we retrieved and matched respective data sets from the Human Protein Atlas (HPA; version 23.0) (Thul *et al*, 2017), Gene Ontology (GO) (Gene Ontology Consortium *et al*, 2023) and UniProt (UniProt Consortium, 2023). The data sets varied largely in the number of terms, the hierarchy between the terms and their reliability scores: HPA provided 40 terms structured in a simple hierarchy with four levels of reliability, the GO ‘cellular component’ subset consisted of 4,058 terms in an ontology graph with 26 evidence codes, and UniProt used 542 subcellular location terms in a simpler ontology graph and four evidence codes. We first mapped and aligned the three public data sets and our data set to a total of ten subcellular location terms: endoplasmic reticulum, Golgi apparatus, plasma membrane, nucleus, lysosome, endosome, mitochondrion, cytosol, cytoskeleton, cell projection (Methods, **EV Fig. 3A**). The RESOLUTE data set included the highest number of annotations with 1,107 defined for 403 SLCs, followed by 741 annotations for 409 SLCs by GO, 469 annotations for 355 SLCs by UniProt, and 339 annotations for 231 SLCs by HPA. We found an overlap of 178 SLCs covered by all four sources, and another 146 SLCs between RESOLUTE, GO and UniProt (**Fig. 3B**).

A considerable amount of these annotations was of weaker confidence, labelled as uncertain, based on computational prediction or based on author/curator statements without experimental evidence. As adding up annotations from different sources increases the false positive rate, we stringently filtered the annotations to experimentally derived, high-confidence localization annotations before merging the data sets (Methods). This high-confidence subset encompassed 434 annotations for 340 SLCs by RESOLUTE, 469 annotations for 300 SLCs by GO, 312 annotations for 247 SLCs by UniProt and 284 annotations for 213 SLCs by HPA (**Fig. 3C**). Merging the four high-confidence subsets, we arrived at 873 localization annotations for 418 SLCs in total, i.e. an average of about 2 locations per SLC. 487 location annotations (55.8%) were supported by a single source, 386 annotations by multiple sources (44.2%), with only 52 annotations (6.0%) supported by all four data sources (**Fig. 3D**). All sources reported Plasma membrane as the most frequent location of SLCs, ranging from close to 100 SLCs in HPA to almost 200 SLCs in GO, and corresponding to almost half or more of the SLCs annotated in the respective sources. Up to seven different locations were annotated for individual SLCs, but the number of locations per SLCs were distributed similarly across all sources (**EV Fig. 3B**).

To judge the level of agreement on the high-confidence annotations between the four data sources, we treated the localization annotation as a multi-label classification problem. We calculated for each pair of data sets the precision and recall per location and combined those metrics to an overall accuracy score across all locations (micro-averaged F1-score (Takahashi *et al*, 2022)). We found the overall highest score and therefore agreement between GO and UniProt annotations. The RESOLUTE data set agreed equally well with both UniProt and GO annotations and to a lower extent with HPA, which generally showed least agreement with all other three annotations (**Fig. 3E**). Comparing the different annotation sources per localization individually, we found the highest agreements for mitochondrion, plasma membrane and Golgi. In contrast, lower agreement was observed for ER location (**EV Fig. 3C**).

Assessing the novelty of high-confidence experimental subcellular location annotations from the RESOLUTE data set, we found for about a third of the SLC family (154 SLCs; 34.5%) previously experimentally derived annotations confirmed. For another third (138 SLCs; 30.9%) RESOLUTE provided experimental evidence for locations in addition to previously reported ones. Most frequent among those were Golgi, Nucleus and Cell projection locations, which might be influenced by the tagged overexpression of the SLC or also by the inherently subjective manual annotation (**EV Fig. 3D**). RESOLUTE reported the first experimental annotations for 48 SLCs (10.8%), including 15 SLCs that contained no prior location annotations at all (**Fig. 3F**). Among those, we found the glucose transporter SLC45A1 and the orphan MFSD11 localized at the lysosome, as well as the orphan transporter NIPAL2 at the Golgi and plasma membrane (**Fig. 3G**).

In the consolidated data set from public sources and our experimental work, more than half of the SLCs (n=262; 58.6%) were annotated at the plasma membrane, but only 70 of them exclusively. The second most frequent exclusive location was mitochondria, with 32 SLCs exclusively annotated and 83 SLCs (18.6%) annotated overall. 155 SLCs (34.7%) had one location and 139 SLCs (31.1%) had two locations annotated. The most common pair of locations was plasma membrane and cell projection (27 SLCs), followed by plasma membrane and Golgi apparatus (22 SLCs). 75 SLCs (16.8%) had three locations and another 49 SLCs (11.0%) had more than three locations annotated. 29 SLCs (6.5%) had no high-confidence location annotation at all (**Fig. 3H**).

### A web portal for integrating data and knowledge on the SLC superfamily

The functional characterization efforts of the RESOLUTE and REsolution consortia created numerous rich and diverse data sets (**Fig. 4A**). To synthesize the different aspects of SLC function into a comprehensive SLC knowledgebase, we integrated our resources and analyses in a database and consolidated selected information from the public domain, such as subcellular localization, structure, tissue expression, disease associations, availability of assays and chemical compounds as well as knowledge from the literature (**Table 1**). We made this SLC-centric data and knowledgebase available via a web portal (https://re-solute.eu) (Methods; **EV Fig. 4**).

**Figure 4.**
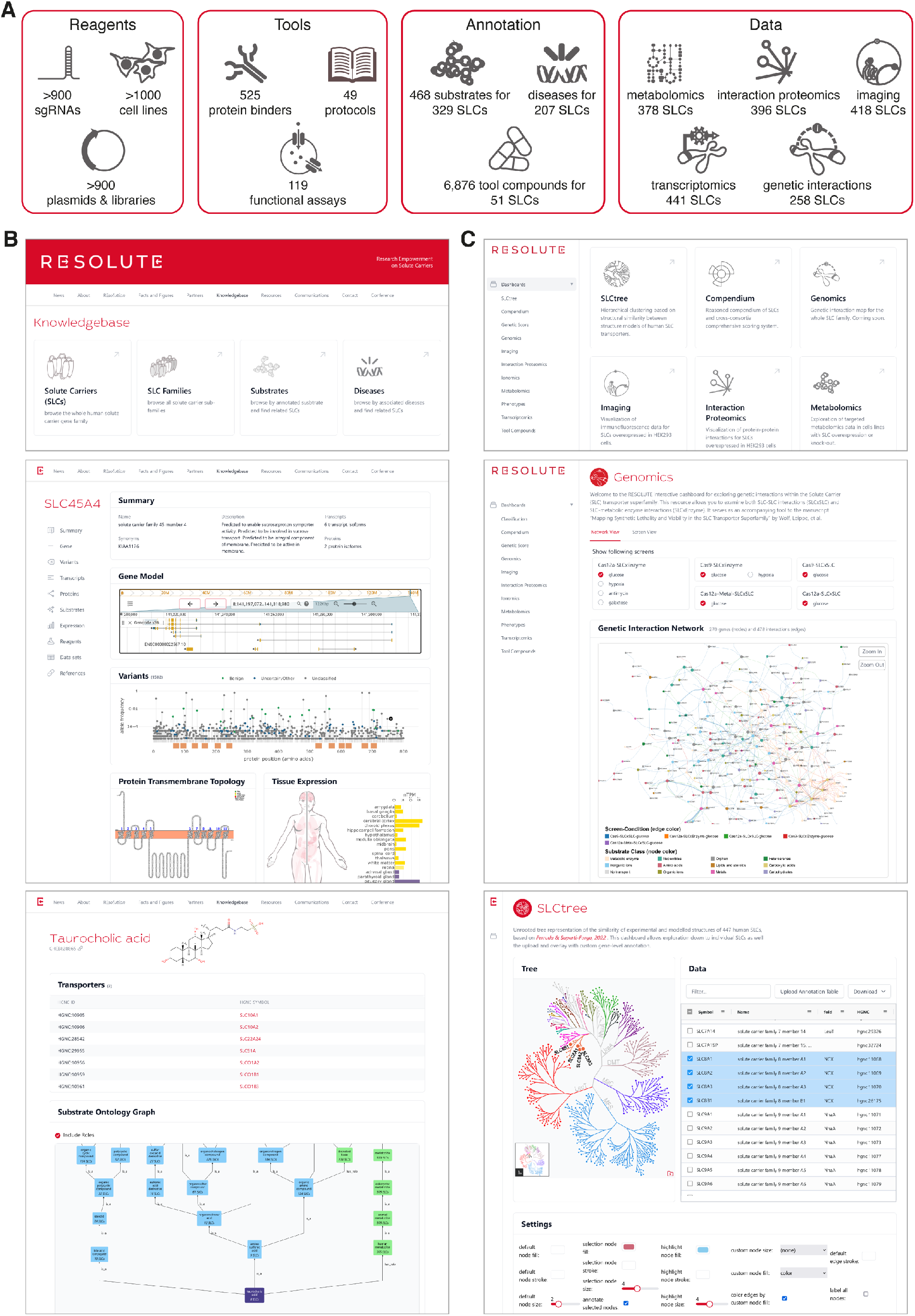
The RESOLUTE web portal. **(A)** Schematic overview of resources generated by RESOLUTE and REsolution consortia. **(B)** Exemplified screenshots of the RESOLUTE knowledgebase, showing the landing page (https://re-solute.eu/knowledgebase) in the upper panel, the detailed view of SLC45A4 in the middle panel and the substrate view for taurocholic acid in the lower panel. **(C)** Exemplified screenshots of the RESOLUTE dashboards, showing the landing page (https://re-solute.eu/resources/dashboards) in the upper panel, the genomics dashboard visualizing the combined genetic interaction network of 5 selected screens in the middle panel and the SLC tree dashboard in the lower panel, demonstrating customization of the unrooted tree representation with colouring by fold and highlighting of the SLC8 family.

**Table 1.** Resources integrated in the SLC data- and knowledgebase.

| resource / reference | version / date | used for | context |
| --- | --- | --- | --- |
| ChEBI | 229 | unification of substrate terms; ontology relationship (generalized querying and analyses); compound properties | knowledgebase / dashboard |
| ClinVar | 2022-02-01 | variant-trait associations | knowledgebase |
| (Digles *et al*, 2024) |  | SLC-specific tool compounds | dashboard |
| EFO |  | unification of disease terms | landscape |
| Ensembl |  | gene model annotations | knowledgebase |
| (Ferrada & Superti-Furga, 2022) |  | fold annotation and structural similarities | knowledgebase / dashboard |
| (Frommelt *et al*, 2024) |  | analysis of the SLC interactome | dashboard |
| Genebass | 0.7.8-alpha | variant-trait associations | knowledgebase |
| gnomAD | 2.1.1 | variant-trait associations | knowledgebase |
| GO |  | localization annotation; unification of disease terms | knowledgebase |
| HGNC | 2024-01-17 | SLC definition; gene-level annotations; identifier matching | knowledgebase |
| Human Protein Atlas | 23.0 | tissue-level gene expression (Uhlén *et al*, 2015);<br>subcellular localization (Thul *et al*, 2017) | knowledgebase |
| IEU OpenGWAS project | 5.9.0 | variant-trait associations | knowledgebase |
| LitVar | 2022-02-01 | variant-trait associations | knowledgebase |
| Mondo | 2024-06-04 | unification of disease terms; ontology relationship<br>(generalized querying and analyses) | knowledgebase /<br>dashboard |
| NCBI | 2023-10-09 | functional summaries; functionally relevant links to<br>literature (GeneRIFs) | knowledgebase |
| Open Targets | 6 | variant-trait associations | knowledgebase |
| Orphanet | 5.52.0 | variant-trait associations | knowledgebase |
| Prioriy Index<br>(Fang *et al*, 2019) |  | variant-trait associations | knowledgebase |
| Protter |  | transmembrane topology visualizations | knowledgebase |
| UniProt | 2024_01 | protein-level annotations; localization annotation; variant-trait associations | knowledgebase |
| (Wiedmer *et al*, 2024) |  | transcriptomics & metabolomics analyses of SLC<br>overexpression cell lines | dashboard |
| (Wolf *et al*, 2024) |  | genetic screens on SLC single and double knockout cell<br>lines | dashboard |
| this study |  | manual curation of SLC substrates; SLC substrate<br>classification; immunofluorescence images of SLC<br>overexpression cell lines; subcellular localization<br>annotation; SLC tree layout | knowledgebase /<br>dashboard |

Different sections enable browsing and querying this unique transporter-focused resource. Next to sections summarizing the aims, mission, and project outputs of the RESOLUTE and REsolution consortia, we provide a comprehensive knowledgebase (https://re-solute.eu/knowledgebase) (**Fig. 4B**). Here, we integrated general knowledge on SLC genes with transporter specific annotations and public data sets. A dedicated page for each SLC member covers basic annotation on the gene-, transcript- and protein-level and matched identifiers. It also includes evidence on gene variants, a visualization of the transmembrane topology, subcellular localization and tissue expression levels, and a summary of experimental data available. The pages also show the results of the curated literature mining for transported substrates and of the data mining for disease associations for each SLC. In addition to the SLC pages, we implemented representations of the compound and disease ontologies which allow finding and grouping SLCs by searching or browsing through the ontology graphs of ChEBI and Mondo, respectively. Both the directly annotated SLCs and the SLCs that were mapped indirectly via annotation to more specific terms can be accessed from every ontology term.

The section on resources (https://re-solute.eu/resources) provides information on and access to reagents, tools, experimental data, and results generated by the consortia. The reagents cover sgRNAs, plasmids and cell lines, and the tools made available to the scientific community are protein binders, functional assays and protocols. Our systematic multi-omics characterization of the generated cell lines led to large-scale experimental data sets and the key insights from these individual data sets were described in accompanying manuscripts (Wiedmer *et al*, 2024; Frommelt *et al*, 2024; Wolf *et al*, 2024). We present the results of our analyses on these data sets using a range of interactive dashboards (**Fig. 4C**). By visual exploration, they enable generation of hypotheses on functional properties of SLCs. A dashboard on the structural classification of SLCs enables custom annotations and visualizations of the SLC tree. Dashboards on the transcriptomics and targeted metabolomics data allow exploration of the regulated transcripts and metabolites, respectively, in each analysed cell line (Wiedmer *et al*, 2024). Dedicated ‘gene’ and ‘compound’ views allow for inspection of the regulation of measured genes and metabolites across all cell lines. The interaction proteomics dashboard enables exploration of bait- or prey-centric interaction networks extracted from the full SLC interactome (Frommelt *et al*, 2024). To expand the context of a network, external protein-protein interaction resources (currently BioGRID and CORUM are supported) can also be overlayed. The genomics dashboard visualizes the integrated genetic interaction network, generated from nine different genetic-interaction screens (Wolf *et al*, 2024). Magnitude and significance of each interaction as well as the effect of the double knockout compared to corresponding individual single knockouts can be analysed at the ‘screen’ view. The imaging dashboard provides access to all immunofluorescence images generated. After selecting a cell line, a stain combination and a view of the corresponding immunofluorescence image can be freely levelled, magnified and downloaded. This dashboard also supports a side-by-side view of two different cell lines or staining. The tool compounds dashboard shows the results of data mining efforts on active compounds for SLCs (Digles *et al*, 2024), and can be used to identify potential tool compounds for an SLC of interest. Finally, the compendium dashboard allows integrated analysis of selected SLC annotations by interactively constructing and navigating a sunburst diagram with an arbitrary number of custom layers and filters. This might lead to interesting insights, e.g. combining fold with subcellular localization and substrate class layers.

To maximize impact of all data generated, we adhered to the FAIR principles (Wilkinson *et al*, 2016). Data sets were released open access using a permissive license. We implemented identifiers, adhered to standardized and open data formats. Meta data, annotations and analysis results are available as structured data or via an API specifically developed for programmatic access. The raw data sets can be downloaded from the web portal. To ensure sustainability, all data sets are also being uploaded to public, domain-specific repositories.

### Construction and application of a multimodal functional landscape of the SLC superfamily

To gain an integrative overview of the functional landscape of human solute carriers, we integrated eight selected, orthogonal data sets on phenotypic profiles of the SLC superfamily. Next to the transcriptional and metabolic profile upon SLC overexpression (Wiedmer *et al*, 2024) we chose the protein interaction network (Frommelt *et al*, 2024), subcellular localization annotation, tertiary structure (Ferrada & Superti-Furga, 2022), the tissue expression pattern (Uhlén *et al*, 2015), substrate annotations as well as disease associations (**Table 2**). On average, 6.9 of those modalities were available per SLC, with a large majority of SLCs (n=404; 90.4%) represented in six or more modalities (**EV Fig. 5A**). The selected modalities encompassed different functional aspects and were represented by fundamentally different data types. For successful integration, we transformed measurements or annotations of each modality to a comparable data type and scale, namely SLC-SLC pair similarity matrices. Briefly, we applied data-type specific distance measures and harmonized the resulting distances to comparable dissimilarities by normalization and transformation (Methods; **EV Fig. 5B**). Correlating the resulting SLC-SLC dissimilarities showed some expected agreements (e.g. structure with substrate, and interaction proteomics with subcellular localization), but the observed correlations are very small (maximum *r_Pearson_*=0.13), indicating that each modality provides additional, orthogonal aspects to SLC function (**EV Fig. 5C**). Merging the modality-specific dissimilarities resulted in an overall similarity for each of the 99,681 possible SLC-SLC pairs (**EV Fig. 5D**). Finally, we employed the UMAP algorithm (McInnes *et al*, 2018) to represent the data manifold underlying the overall similarity matrix as a graph, which was then embedded in a two-dimensional SLC landscape (**Fig. 5A**).

**Figure 5.**
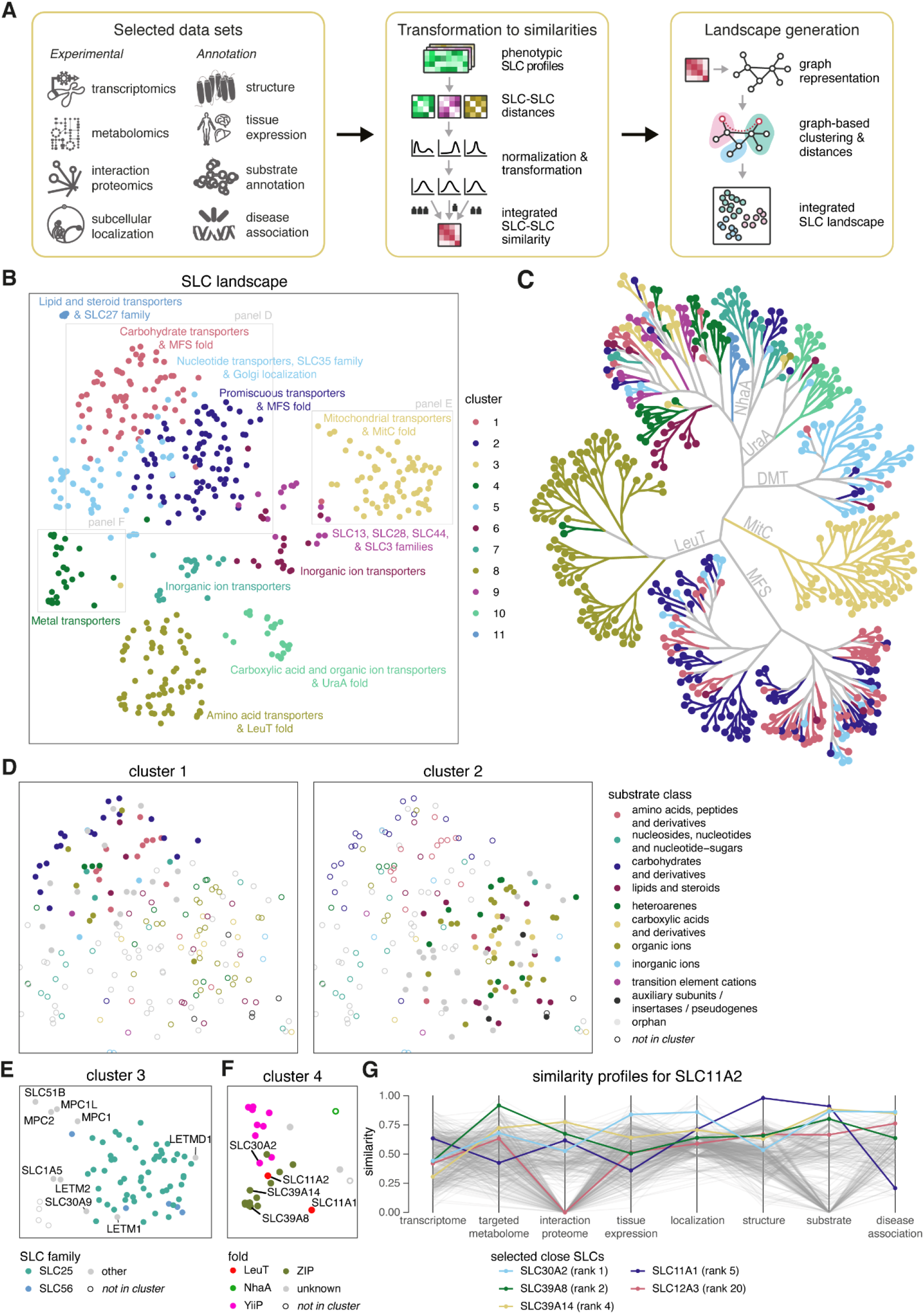
An integrated, functional landscape of the human SLC superfamily. **(A)** Schematic overview of the workflow generating the SLC landscape. The eight selected source data sets are illustrated in the first box. The second box details the transformation of source data to integrated SLC-SLC similarities. And the third box indicates how the UMAP algorithm was used to derive a neighbourhood graph and the final landscape. **(B)** The two-dimensional representation of the functional landscape of 447 SLCs. Results of the high-dimensional graph-based clustering are indicated by colour and labelled by simplified but expressive cluster names. Selected areas from panels D, E and F are indicated by grey boxes. **(C)** The cluster membership of each SLC is visualized on the structure-based SLC tree, using the same colour scheme as in panel B. The six largest folds are labelled on their respective branch. **(D)** Cut-out of the functional SLC landscape highlighting the clusters 1 and 2 and demonstrating the different predominance of substrate classes. **(E)** Cut-out of the functional SLC landscape highlighting the cluster 3 and demonstrating the overrepresentation of the mitochondrial SLC25 family members. The mitochondrial zinc transporter SLC30A9 is highlighted. **(F)** Cut-out of the functional SLC landscape highlighting the cluster 4 and demonstrating the two major folds of metal transporters (YiiP and ZIP). SLC11A2, featuring a diverging LeuT fold, and some of its neighbours are highlighted. **(G)** Profile plots of the similarity of all SLCs to SLC11A2, in each of the eight modalities that were used in constructing the functional landscape. Selected SLCs, which are close on the landscape, are highlighted, among them the two manganese transporters SLC39A8 and SLC39A14.

**Table 2.** Data sets integrated in the functional SLC landscape.

| data set | source | SLCs covered | data type | distance measure | integration weight |
| --- | --- | --- | --- | --- | --- |
| transcriptomics | RESOLUTE (Wiedmer *et al*, 2024) | 441 | profiles of differential transcript abundances | Euclidean | 3 |
| targeted metabolomics | RESOLUTE (Wiedmer *et al*, 2024) | 378 | profiles of differential metabolite abundances | Euclidean | 3 |
| interaction proteomics | RESOLUTE (Frommelt *et al*, 2024) | 396 | sets of interacting proteins | co-occurrence of interactors | 3 |
| subcellular localization | RESOLUTE (this study) | 418 | annotated lists of cellular compartments | term co-occurrence | 1 |
| structure | (Ferrada & Superti-Furga, 2022) | 447 | atomic coordinates | structural alignments | 3 |
| tissue expression | Human Protein Atlas (Uhlén *et al*, 2015) | 443 | profiles of gene expression in tissues | correlation-based | 1 |
| substrate annotation | this study | 329 | annotated lists of chemical compounds | chemical fingerprints | 2 |
| disease association | REsolution (this study) | 226 | annotated lists of disease associations | semantic similarity | 1 |

Clustering the underlying high-dimensional graph identified 11 distinct communities of SLCs on the landscape (**Fig. 5B**). Overall, the clusters recapitulated some of the major folds (**Fig. 5C**), for example, cluster 8 included all SLCs featuring a LeuT fold, except for the two SLC11 family members that located in cluster 4 together with other metal transporters. Other folds were spread across two or more clusters such as the MFS fold that had 54 and 66 of its 138 members in clusters 1 and 2, respectively (**Fig. 5D**). The separation of MFS fold members appeared to follow substrate specificity, as 30 out of the 42 MFS fold members with known substrates in cluster 1 were classified as amino acid or carbohydrate transporters, compared to only 3 out of 47 MFS fold members with known substrate in cluster 2. In general, the substrate specificity in cluster 2 appeared to be much more promiscuous.

The quality and coherence of the clusters was assessed via their mean silhouette scores (Rousseeuw, 1987), which were positive for all clusters and ranged from 0.08 for the more loosely defined cluster 1, to 0.73 for the very coherent cluster 11 (**EV Fig. 5E**). To study the clustering more closely, we performed a mutual information analysis systematically comparing the clusters on the landscape with a defined set of discrete SLC properties, namely fold, family, substrate class, subcellular localization and number of modalities. While fold annotation showed highest coherence with landscape clusters, the other properties also had considerable mutual information with the overall clustering except for the number of modalities, which did not contribute to clustering (**EV Fig. 5F**). A detailed enrichment analysis of all the SLC property levels for each cluster showed between 2 and 10 significantly overrepresented property levels for each cluster (**EV Fig. 5G**).

An example of the contribution of subcellular location to the clustering can be found in cluster 3, resembling the mitochondrial localized solute carriers (**Fig. 5E**). While it is dominated by members of the MitC fold members, it also includes all sideroflexins featuring the rather distant SLC56 fold. Another interesting member of cluster 3 is SLC30A9, which is a zinc transporter featuring a YiiP fold. On the SLC landscape, SLC30A9 is relatively dissimilar to the other YiiP fold members, which are all in cluster 4. Indeed, we found SLC30A9 exclusively localized at the mitochondria, supporting previous computational (Kowalczyk *et al*, 2021) and experimental evidence (Rensvold *et al*, 2022).

Cluster 4 covered 25 of the 31 metal transporters. Here, substrate class appeared to be the major determinant as the cluster integrated the two well known, but rather distinct YiiP and ZIP folds, corresponding to SLC30 and SLC39 family members, respectively (Wiuf *et al*, 2022; Pasquadibisceglie *et al*, 2022; Zhang *et al*, 2023). Next to those two typical families for metal transport, the landscape also positioned SLC11A2 – also known as Divalent Metal Transporter 1 (DMT-1) – at the center of this cluster (**Fig. 5F**). In particular, the manganese transporters SLC39A8 and SLC39A14 were two of the four closest SLCs to SLC11A2. Inspecting the similarities on each of the eight modalities that were used to construct the landscape, revealed high similarities in targeted metabolome profiles, interaction networks, annotated substrates and associated diseases (**Fig. 5G**). Indeed, previous studies have shown that all three proteins can transport the same divalent cations (Polesel *et al*, 2023; Garrick *et al*, 2006). In addition, manganese and iron homeostasis is dysregulated in various animal models when transporter function is disrupted (Fujishiro *et al*, 2012; Tuschl *et al*, 2013; Chen *et al*, 2018). There is also evidence that altering SLC39A14 expression levels influences the expression of SLC39A8 and SLC11A2 in different mouse organs, further suggesting that these transporters are functionally related (Xin *et al*, 2017). Together, these findings support the remarkably close positioning of these three transporters on the functional SLC landscape, regardless of their different structural folds.

In summary, the SLC functional landscape seems to represent a useful integrative and comparative representation of properties of this rather understudied supergroup of membrane transporters and, as tool, offers insights beyond those that can be gained by manual annotation of the different properties of individual transporters.

## Discussion

The complexity of the task of assigning functional properties to a large group of gene products that are related only by a shared principle, that of transporting inorganic and organic molecules across lipid bilayers without hydrolysing ATP, is enormous (César-Razquin *et al*, 2015; Superti-Furga *et al*, 2020; Hediger *et al*, 2004). Human membrane transporters seldom transport only one chemotype and functional redundancy is built-in in circuits fundamental for all cellular functions such as provision of energy and macromolecular building blocks, control of ionic strength and osmolarity for protein folding and chemical reactions, control of pH, redox and membrane potential. In retrospect, it is difficult to think of another class of proteins whose function is as broadly intertwined with basic biophysical effects as SLC transporters. In comparison, GPCRs, transcription factors, proteases, and RNA binding proteins all may have very large individual effects but are all essentially conveying signals. Transporter activity however changes the very chemical makeup of the environment the proteins operate in. Thus, direct functional consequences in the systematic interrogation of SLCs are obscured by a plethora of compensatory mechanisms and hinder the unambiguous assignment of transporter substrates and functions.

Despite these inherent difficulties, we have been able to assign functional properties to each SLC, predominantly defined by the integration of measurements upon elevated protein expression. In total, we used dimensions derived from eight different modalities to calculate an integrated virtual landscape in which each SLC finds a specific location based on its corresponding data and annotation. Although this may not always allow derivation of a definitive or unequivocal functional assignment to each SLC superfamily member, it allows for a meaningful comparison of neighbourhoods and relative multidimensional distances of properties between SLCs. The molecular profiles employed in this study (transcripts, metabolites, interacting proteins), representing measurements derived from the genetic control of protein expression under comparable, controlled conditions, were weighed more strongly when deriving the functional landscape. We think that this is where the power of the approach lies and was not possible, for the first two dimensions, before the advent of CRISPR/Cas technology (Bock *et al*, 2022; Wang & Doudna, 2023). It is common to find sceptics of large-scale biology arguing that only in-depth dedicated studies allow for real new biological insights. While parallel approaches cannot replace dedicated studies, it is precisely the comparative nature of parallel assessment under controlled conditions that allows for objective conclusions (Hieter & Boguski, 1997; Ideker *et al*, 2001; Collins *et al*, 2003). Until now, most insights on SLC protein function addressed only one or a few members of an SLC family or of SLCs acting in concerted action. It is the global, integrative and comparative nature of the SLC landscape that allows for the identification of accrued properties not inferable by subfamily membership alone.

The value of the experimental data obtained in this concerted effort is potentially limited by several factors mentioned in the individual accompanying manuscripts (Wiedmer *et al*, 2024; Frommelt *et al*, 2024; Wolf *et al*, 2024). One limitation that is apparent also in the experimental data set for subcellular localization described here, is that protein overexpression models may lead to protein accumulation in the ER and therefore an overrepresentation of annotations to this location or connected compartments such as nucleus and Golgi, which is reflected in the proportionally large number of novel assignments to these compartments by the RESOLUTE data set. However, for systematic experimental subcellular localization, it is necessary to use overexpression models due to the lack of specific antibodies, in particular for SLCs (Gelová *et al*, 2024; Ayoubi *et al*, 2023). This limitation is also apparent from the observation that HPA, consisting of antibody-based evidence of subcellular localization, showed least agreement with other annotation sources. Importantly, the SLC superfamily-wide localization data presented in this study provides first experimental evidence for 48 SLCs. Overall, it is a valuable resource for understanding SLC function as well as for assay design and drug discovery, for which knowledge of subcellular localization is crucial (Dvorak *et al*, 2021).

To partly correct for specific bias of the experimental approaches, we include data not likely to be affected by similar bias, such as protein structure analysis, chemical substances known to be transported by individual SLCs and genetic association with diseases. While the protein structure analysis using AlphaFold2 predictions was published recently (Ferrada & Superti-Furga, 2022), the substrate annotation and disease associations required systematic collection, annotation and curation. Compared to the initial annotation of SLC substrates a few years back (Meixner *et al*, 2020), our integrative analysis required stronger experimental evidence for substrates. Despite the more stringent criteria applied, the fact that research on transporters was gaining traction over the last years led to a higher number of SLCs with known substrates. However, substrate annotation is far from complete. Discovery of the full panel of substrates might be partially hindered by the common use of gene/protein names that imply a particular function as a given fact, thereby demotivating research efforts on more physiologically relevant function or yet unknown substrates. For example, SLC2A1 (currently named ‘solute carrier family 2 member 1’) had a previous name ‘human T-cell leukemia virus (I and II) receptor’. In this sense, the use of the HUGO nomenclature for SLC genes and proteins leads to less bias than the one induced by historically adopted, imprecise names. Moreover, the limitation of any literature-based annotation remains the collective bias of research topics towards well studied areas, resulting in understudied genes (Edwards *et al*, 2011).

A comprehensive overview of pathogenic genetic alterations in SLC genes has not been available since genetic studies are typically focused on individual SLCs and specific diseases (Li *et al*, 2021), families (Kölz *et al*, 2021), refer only to a limited population (Rajman *et al*, 2020; Mir *et al*, 2022) or contain only limited genetically-encoded aspects (Schaller & Lauschke, 2019). We undertook a systematic data mining and curation process for all SLCs, consolidating previously fragmented information. The curated data set represents a unique resource of genetic, clinical, and experimental evidence, and enabled the inclusion of understudied SLCs, which have not yet been clinically recognized, revealing translational implications for mutations observed in different SLC families and individual SLCs. For instance, the SLC44 and SLC39 families were characterized by a high number of likely pathogenic associations. While many individual transporters within these families have been reported in various contexts, the two main drivers were SLC44A2 and SLC39A8. For SLC44A2, encoding a choline transporter, ClinVar reports no pathogenic variants but recent studies identified its role in a range of disorders, such as the regulation of thrombosis and hearing impairment (Bennett *et al*, 2020; Koehl *et al*, 2023). On the other hand, SLC39A8, encoding a metal ion transporter, has emerged as one of the most pleiotropic genes identified in GWAS studies (Pickrell *et al*, 2016). Its over 40 distinct associated disorders extend beyond current clinical applications (Park *et al*, 2015; Choi *et al*, 2018), some of which have only recently been subject to in-depth investigation (Nebert & Liu, 2019; Haller *et al*, 2018; Nakata *et al*, 2020; Costas, 2018). One limitation of the human genetics data set are potential biases, such as population-based variations and sample size constraints inherent in the mined databases (Fitipaldi & Franks, 2023). These biases may skew the representation of certain genetic variants, particularly when certain populations are not well represented (Schaller & Lauschke, 2019), making it difficult to generalize findings across diverse populations. Another challenge arises from the nature of GWAS, where individual associations may result in false positives due to linkage disequilibrium (Slatkin, 2008).

To empower future research on solute carriers, we set up a web portal allowing the scientific community to access and explore the resources, tools, analyses and data sets collected and generated by the RESOLUTE and REsolution consortia. Addressing the typical limitations of such a knowledgebase, we chose a modern software architecture that minimizes maintenance efforts and updateable modular workflows on flexible data models to counteract outdated information. Data sets and resources are also transferred to public, domain-specific repositories for increased sustainability.

The data- and annotation-informed integration to a global SLC landscape allowed us to define functional distances between SLCs and group them into 11 clusters, each resembling a mix of different known properties of SLCs. Next to investigating an SLC of interest via the transporters featuring the most similar phenotypic profiles, the integrated landscape allowed extension of this guilt-by-association principle also to the functionally closest genes, combining local and global features of the integrated data sets. Examining the landscape enabled us to uncover interesting relations between SLC family members. We found the iron transporter SLC11A2 located close to the manganese transporters SLC39A8 and SLC39A14. The similarity between these three transporters could be beneficial for designing novel therapeutics. Loss of function mutations in SLC39A8, SLC39A14, or SLC11A2 lead to three distinct clinical presentations. Bi-allelic mutations in either SLC39A8 or SLC39A14 transporter lead to the rare diseases Disorder of Glycosylation type IIn (CDG2N; LOF mutations in SLC39A8) or Hypermanganesaemia with Dystonia 2 (HMNDYT2; LOF mutations in SLC39A14), both characterized by abnormal blood Mn levels (Tuschl *et al*, 2013; Winslow *et al*, 2020; Anagianni & Tuschl, 2019). Conversely, SLC11A2 loss of function or reduced expression results in microcytic anemia (Mims *et al*, 2005; Beaumont *et al*, 2006; Shawki *et al*, 2015). The different clinical phenotypes may result from differences in Mn or Fe binding affinity or total transporter abundance *in vivo*; therefore, altering the abundance of a similar transporter may restore metal homeostasis. For example, pharmacologically increasing gene expression of SLC39A8 in the duodenums of patients encoding SLC11A2 loss of function mutations could increase Fe levels in this organ, since both proteins mediate metal transport across apical membranes of polarized cells (Gunshin *et al*, 1997; He *et al*, 2006), and SLC39A8 expression leads to increased iron uptake in HEK293 cells (Wang *et al*, 2012). Overall, this example demonstrates that the similarity between transporters can be used to generate therapeutic strategies for SLC-related disorders.

Beyond the few examples highlighted in this study, the landscape offers much more to explore. While this functional allocation is only a starting point for future work, it provides the groundwork for real functional implications of hundreds of SLCs that were uncharted before and multiplies the collected previous knowledge on the subject. A limitation due to the simplification of a two-dimensional representation of the landscape is that the closeness between two transporters is not always reflected in vicinity on the depicted map of the landscape. Visual interpretability may profit from novel visualization techniques (Hütter *et al*, 2022). Variations of the landscape could be generated by tuning the weights of the different integrated data sets or by the inclusion of additional dimensions. Transforming the selected data sets to SLC-SLC similarities allowed for more straightforward integration but also sacrificed potentially valuable information. Integration via prior-knowledge networks could lead to successful integration while preserving most information, and at the same time open up interesting applications of machine learning (Fortelny & Bock, 2020). While not used in this study, prior-knowledge networks have been used at a smaller scale by employing the Reactome pathway database for linking gene expression to metabolite abundances (Wiedmer *et al*, 2024; Milacic *et al*, 2024). A promising route will be the integration of our comprehensive data sets on human solute carriers to knowledge graph initiatives, such as the recently published BioCypher (Lobentanzer *et al*, 2023).

When we confronted the community with the necessity to mount a concerted action on human membrane transporters ten years ago, particularly on the rather many and partly obscure solute carriers, the knowledge landscape was different (César-Razquin *et al*, 2015). Less than a handful of experimentally determined structures of human SLCs were available and the main functional assay consisted of injecting human mRNA encoding an SLC in frog oocytes and measuring the differential accumulation of a radiometrically labelled suspected substrate. Given the importance of the interface between the chemical and the biochemical world, and the fact that it is mainly governed by transporters, a call for action seemed appropriate. Where do we stand ten years after? The research landscape on SLCs and related transporters has changed dramatically, such that the SLC family recently lost its title of the ‘most asymmetrically studied gene family’ to the family of pleckstrin homology domain containing proteins (**Fig. 6**). There is a new level of awareness and engagement in SLC research as exemplified in some spectacular recent studies (Wang *et al*, 2021; Morioka *et al*, 2018; Luongo *et al*, 2020; Adelmann *et al*, 2020; Liu *et al*, 2023), the ability to chemically address the target class has improved dramatically (Wang *et al*, 2020; Galetin *et al*, 2024; Dvorak & Superti-Furga, 2023) and there has been an explosion in the elucidation of three-dimensional structures via cryo-EM that has not only clarified transport mechanisms but also revealed the mechanism of action of important therapeutics (Parker *et al*, 2021; Parker & Newstead, 2017; Coleman *et al*, 2019; Côté *et al*, 2011; Srivastava *et al*, 2024). Moreover, a wider role of SLCs in drug uptake has been recognised (Girardi *et al*, 2020). There are many more functional assays than a decade ago, especially involving intact human cells.

**Figure 6.**
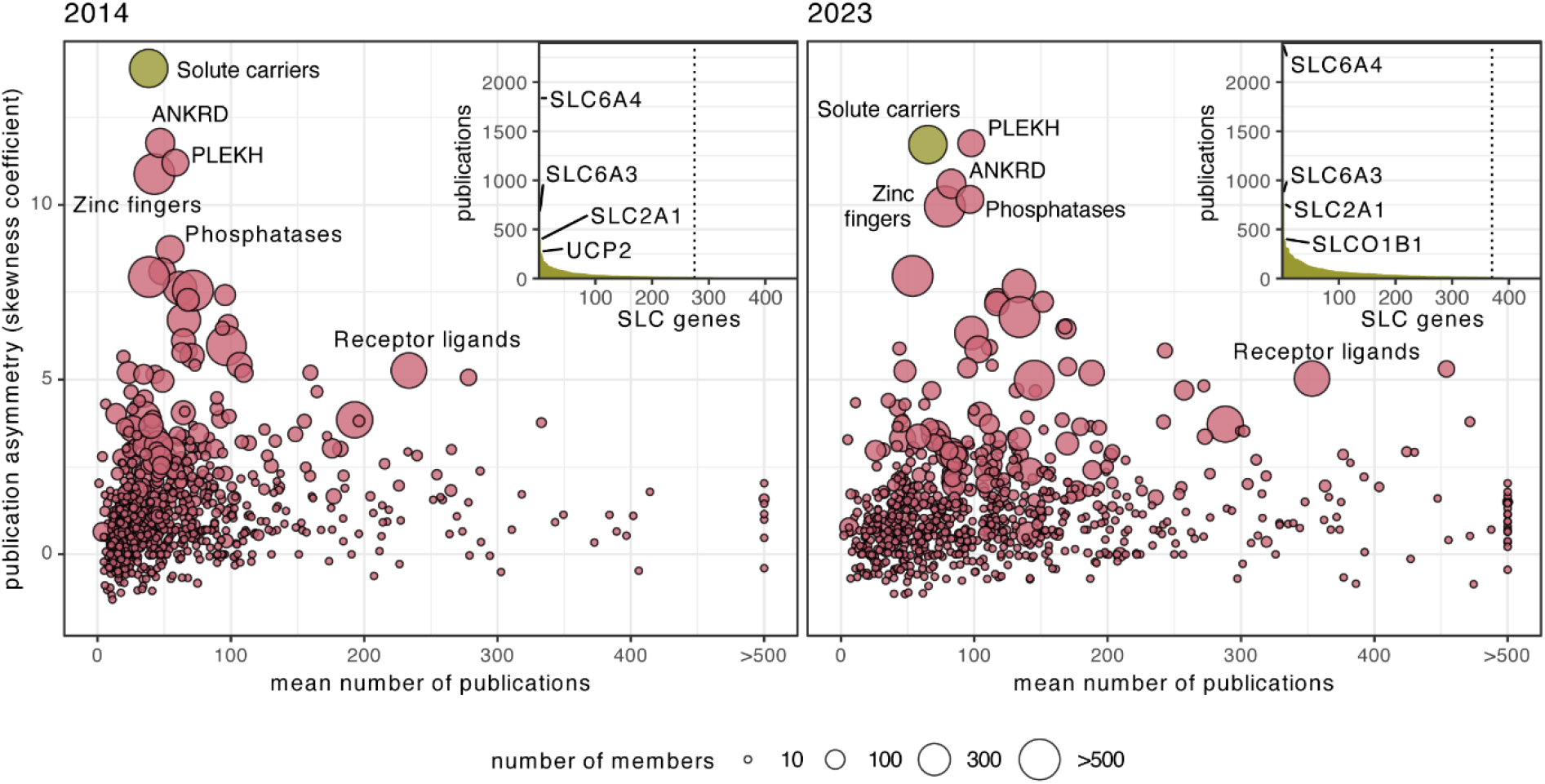
Evolution of publication asymmetry over the last 10 years. Publication asymmetry of gene families compared to their members’ mean number of publications, as shown in (César-Razquin *et al*, 2015). Publications were considered up to the end of 2014 on the left panel and up to the end of 2023 on the right panel. Over the last decade, publication asymmetry increased for some families, such as the group of pleckstrin homology domain containing (PLEKH) genes or the group of phosphatases, it remained constant for families such as the group of receptor ligands and decreased for other families, particularly for the solute carrier superfamily. Point size corresponds to the size of the gene family and selected gene families are labelled. The inset ranks the SLC genes by their number of associated publications, demonstrating the asymmetry within this superfamily. The four most studied SLCs are indicated. The dashed line marks the threshold to SLCs with less than 15 associated publications, corresponding to 181 SLCs as of 2014 and 86 SLCs as of 2023. Publication counts per gene are based on the ‘gene2pubmed’ and ‘generifs_basic’ files provided and curated by NCBI. Gene groups as defined by HGNC. Only protein-coding genes were considered. Asymmetry is measured for each group of genes by calculating the skewness (‘moments’ library, version 0.14.1) of the distribution of the number of publications for all genes within the group. A high skewness coefficient indicates a distribution where a few genes in the family concentrate a much higher number of publications than the rest.

The knowledgebase we present here, together with the newly-generated toolbox of assays (Dvorak *et al*, 2021; Digles *et al*, 2024), protein binders (Gelová *et al*, 2024), plasmids and cell lines (Wiedmer *et al*, 2024), and omics data sets (Wiedmer *et al*, 2024; Frommelt *et al*, 2024; Wolf *et al*, 2024) should represent a turning point in the ability of the community to work on this superfamily of proteins. We hope that a publication asymmetry continues to decrease (**Fig. 6**) manifested by an acceleration in the process of creating a more even engagement of the research community to the SLC superfamily. The threshold should be lowered as no SLC is now likely to be completely void of functional suggestions, genetic interactions, protein interaction partners, genetic variants associated with human diseases, structural model, or subcellular localisation. We are convinced that the unusually large effort that was mounted in the last years represents a formidable starting point for research on the largest group of membrane transporters encoded in the human genome and a potential blueprint for similar efforts on other group of neglected human gene products.

## Material and Methods

### Constructing the SLC tree

Pairwise structural similarities were obtained from (Ferrada & Superti-Furga, 2022). Distances were calculated by subtracting similarities from 1 and used for hierarchical clustering using Ward’s criterion (Ward, 1963). Branch lengths of the resulting dendrogram were square-root scaled and visualized as unrooted tree using the equal daylight algorithm of ‘ggraph’ library (version 2.2.1). The tree was further semi-automatically adjusted using Gephi software (version 0.10.1) (Bastian *et al*, 2009).

### Substrate annotation, ontology mapping and substrate class definitions

A team of 16 people in the Superti-Furga group (graduate students and postdoctoral fellows) scanned the primary literature on human solute carriers. The criteria for annotating a compound as a substrate are more strict than in our previous annotation effort (Meixner *et al*, 2020). Only reports with proof from a transport assay using human solute carrier proteins were considered, no homology-based inferences were allowed. Collected compound names were manually mapped to ChEBI terms (Hastings *et al*, 2016). Biologically corresponding terms (tautomers, conjugated bases/acids) were identified and harmonized. For mapped generic terms, which describe not a single compound but rather a class of compounds in ChEBI, the primary literature was re-checked trying to find and match a more specific chemical compound. After a number of iterations, we arrived at 2,044 SLC-compound-publication pairs. Every SLC-substrate annotation has therefore at least one reference to a primary resource, mainly via PubMed identifier.

We defined a set of 9 biologically and biochemically relevant substrate classes based on manual selection of higher-level ChEBI terms (listed in parentheses): ‘lipids and steroids’ (CHEBI:18059, CHEBI:35341), ‘nucleosides, nucleotides and nucleotide-sugars’ (CHEBI:33838, CHEBI:36976, CHEBI:25609), ‘amino acids, peptides and derivatives’ (CHEBI:33709, CHEBI:16670, CHEBI:22860, CHEBI:37793, CHEBI:63534), ‘carbohydrates and derivatives’ (CHEBI:78616, CHEBI:24848), ‘heteroarenes’ (CHEBI:33833, CHEBI:17015), ‘carboxylic acids and derivatives’ (CHEBI:33575, CHEBI:37622), ‘transition element cations’ (CHEBI:33515), ‘organic ions’ (CHEBI:25699), ‘inorganic ions’ (CHEBI:36914). Annotated substrate terms were then matched to those terms, walking down the ontology tree on ‘is a’, ‘is conjugate base of’, ‘is conjugate acid of’, ‘is tautomer of’, and ‘has role’ relations. Then we summarized the classification of substrate terms per SLC. Here an SLC often matched to different classes, either due to multiple annotated substrates matching to different classes, or also due to a single annotated substrate matching to multiple classes. Cases of ambiguous class matching were automatically resolved using the first match based on the order of the classes listed above, and some SLC classifications were manually curated afterwards.

### Disease association mapping

For all variants collected, we assessed the consequences on the canonical transcripts of all SLC genes as annotated in gnomAD via Sequence Ontology (version 2.5.3) (Eilbeck *et al*, 2005) terms. Variants were then filtered for consequences from the ‘protein_altering_variant’ (SO:0001818) sub-ontology graph.

Our workflow to map all collected traits to selected ontologies was relying on three publicly available resources. The first was the Python module OnToma, which processed identifiers from other ontologies and also free text labels to return a corresponding EFO term. For identifiers we considered the first ranked successful match, for free text labels, the process included searching for an exact name match from the EFO OT slim OWL file, an exact synonym match from the OWL file, a mapping from a manual string-to-ontology database, and a high-confidence mapping from EBI’s ZOOMA tool with default parameters. ZOOMA leverages its manually curated annotations from publicly available databases to find potential matches, accompanied by confidence scores. For traits that remained unmapped by OnToma, ZOOMA was used with more lenient confidence thresholds as the second step of the mapping process. The outcomes were manually reviewed to eliminate obvious inaccuracies. Traits that were still unmapped after this secondary step were considered not related to biological associations. The final step involved using EBI’s OxO (Ontology Cross-Reference Service), developed to facilitate the identification of cross-references and synonyms across different ontologies, vocabularies, and coding standards. Our aim was to identify disease traits mapped to EFO or HP terms, as opposed to those mapped to Mondo. This allowed the separation between ‘pathogenic’ traits, associated with terms from Mondo, and ‘non-pathogenic’ traits, associated with terms from either EFO or HP. There were three rather unspecific Mondo terms, which we manually excluded from the ‘pathogenic’ set: ‘alcohol-induced mental disorder’ (MONDO:0002326), ‘generalized anxiety disorder’ (MONDO:0001942), and ‘Mendelian disease’ (MONDO:0003847).

Disease areas were defined as the 37 direct child terms of the ‘human disease’ term in Mondo (MONDO:0700096). A disease association was counted for a specific disease area if it mapped to the disease term or any more specific term in the ontology. Due to the ontology structure, a single disease term may be categorized under multiple disease areas.

### Immunofluorescence imaging

Jump-In T-REx HEK293 cells were generated and cultured as described in (Wiedmer *et al*, 2024) and in detail here: https://zenodo.org/records/7457221 and https://zenodo.org/records/5566805. The detailed protocol for immunofluorescence imaging is available at https://zenodo.org/records/7457346. In brief, cells were seeded on laminin-coated COC high content imaging plates at seeding densities adjusted to yield 80% confluency at the beginning of the staining procedure. After 2 days for attachment and proliferation, cells were treated with 1 µg/ml doxycycline to induce transgene expression for 22 hours or were left untreated. Doxycycline was washed out and cells were grown for another 18 hours. Antibodies or chemical probes were applied to stain 9 cellular compartments and the SLC-HA-tag either prior or post fixation and permeabilization according to manufacturer’s recommendations. Cells were incubated with Mitotracker Orange CMTMRos for 30 minutes, prior to fixation with 3.5% PFA for 25 minutes and washing. Afterwards, samples were permeabilized and blocked with Triton-X, Digitonin and FBS for 30 minutes, followed by washing. Samples were incubated with primary antibodies for 2 hours at 37 °C (Hoechst 33342, HCS Cellmask DeepRed, anti-KDEL, anti-Giantin, anti-LAMP2, anti-PMP70, anti-RAB9A, anti-Na-K-ATPase, anti-Tubulin, anti-HA-tag), washed, incubated with fluorophore-labelled secondary antibodies for 90 minutes at 37 °C and washed again. Plates were imaged with the Yokogawa CV7000 using a 60x water immersion objective. Images were assessed visually by scaling manually to comparable intensities and attributing scores according to the intensity per individual compartment over the total intensity and assigning a confidence score from 1 to 5, with 5 as highest confidence level. Throughout the process of creating a machine learning-based prediction tool for localization of a given protein in HEK293 cells (Baranowski *et al*, in preparation), repeated training iterations of machine learning and the study of mismatches between human annotation and prediction led to several revisions and a thoroughly curated SLC localization annotation list.

### Mapping of subcellular localization annotations

Immunocytochemistry-based annotation of subcellular localizations were obtained from the Human Protein Atlas (HPA; version 23.0) (Thul *et al*, 2017). From the UniProt database (version 2024_01), we obtained Gene Ontology (GO) based subcellular location annotations and the curated set of UniProt annotations. Based on the manual annotation of the RESOLUTE immunofluorescence data set, we selected 10 matching subcellular location terms from the Gene Ontology and identified matching terms in HPA and UniProt. The selected terms were automatically integrated with annotations for any of their sub-terms, resulting in consideration of 1,823 terms in GO, 158 terms in UniProt, and 22 terms in HPA. There was no matching term for cell projection in HPA. We ranked the reliability scores and evidence codes provided by the different annotation sources and considered only the annotation with the highest score in cases of multiple annotations for an SLC to a selected location term within the same annotation source (**EV Fig. 3A**). We then applied a strict filtering to define the high-confidence subcellular localization annotations: for RESOLUTE annotations the confidence score had to be 5 and the signal proportion had to be at least 0.2; UniProt annotations had to have the evidence tag ‘experimental evidence used in manual assertion’ (ECO:0000269); GO annotations had to have the evidence code ‘inferred from direct assay (IDA)’, ‘inferred from mutant phenotype (IMP)’ or ‘inferred from high throughput direct assay (HDA)’; and HPA annotations had to have a reliability score of ‘Enhanced’, ‘Supported’ or ‘Approved’ (**Fig. 3C**).

### Implementation of an SLC-centric web portal

The individual components of the web portal (**EV Fig. 4**) were developed in a GitLab environment (version 14.9.0) and deployed containerized (Docker version 24.0.2).

A relational database in a PostgreSQL system (version 15) served as the data layer for the web portal. To populate the database, we employed TypeScript or Python to develop ETL (extract, transform, load) pipelines for processing result tables of our omics data analyses or public data. Pipelines integrating publicly available data were operating directly on APIs or on downloadable snapshot or source files. Prisma (version 5.5.2) served as an object-relational-mapper, streamlining the implementation between data layer and ETL pipelines.

The web server was implemented in Node.js (version 20) using the Express framework (version 4.17.1). An application programming interface (API) was implemented via PostGraphile (version 4.13.0), providing machine-readable access to the data using the GraphQL query language (https://re-solute.eu/graphiql).

The frontend was implemented in TypeScript using the component-based framework React (version 18.2.0) and was served by an nginx server (version 1.24.0). The GraphQL client urql (version 3.0.3) was handling communication to the backend. Several libraries were employed to implement the user interface and enhance the user experience, in particular Plotly (version 2.27.0) for interactive plots and charts, sigma.js (version 3.0.0) for interactive graphs and networks, AG Grid (version 26.1.0) for customizable data grids, and Tailwind (version 3.3.2) for general styling of components.

Not all data was available in the relational database and therefore accessible via the API. Some data sets were served from the web server in structured data files (JSON format), while all larger data sets (in particular omics raw files) were available form the Microsoft Azure cloud storage.

### Calculating an SLC-SLC similarity matrix

Integration of individual data modalities was achieved by transforming each data set to a similarity matrix. For each modality, we first calculated pairwise distances per SLC-SLC combination (**EV Fig. 5B**). For transcriptomics and metabolomics, the distances corresponded to the Euclidean distance between standard-normalized profiles of logarithmized fold-changes of measured transcripts and metabolites upon SLC overexpression, respectively (Wiedmer *et al*, 2024). For interaction proteomics, the distance was the Jaccard distance between the sets of identified prey proteins, as detailed in (Frommelt *et al*, 2024). For tissue expression, the distance was based on the Spearman correlation 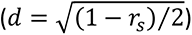 of RNA expression profiles across 50 tissues in the Human Protein Atlas ‘RNA consensus tissue gene’ data (from proteinatlas.org; version 23.0) (Uhlén *et al*, 2015). For subcellular localization, we first determined the pairwise distances of the subcellular location terms based on their overlap of annotated SLCs (Jaccard distance) and then integrated the distances of potentially multiple localizations per SLC-SLC pair to a single distance using the ‘best-match average’ (BMA) method (Gu, 2024). For structure, we employed distances from a recent structural analysis of the solute carrier superfamily (Ferrada & Superti-Furga, 2022) which are based on the overlap of pairwise alignments of experimental or modelled protein structures. For substrate annotation, we calculated the distance between molecular substrates as the Tanimoto coefficient of OpenBabel FP4 fingerprints using the ‘ChemmineR’ library (version 3.54.0) (Cao *et al*, 2008) on SDF format representations of the molecular compounds downloaded from ChEBI (version 22.9). The distance between elemental substrates was defined as Euclidean distance of the scaled ‘Oliynyk’ representation in the ‘ElementEmbeddings’ library (Onwuli *et al*, 2023). Elemental substrate distances were further scaled to a maximum of 1 and then combined with molecular substrate distances to an overall substrate distance matrix, using the maximum distance (i.e. *d*=1) between molecular and elemental substrates. Again, the distances of potentially multiple substrates per SLC-SLC pair were integrated to a single distance using the BMA method. For disease associations, the semantic distance of two disease terms was determined from the graph representation in the Mondo ontology using the ‘Leacock’ distance (Leacock & Chodorow, 1998) implemented in the ‘simona’ library (version 1.0.10) (Gu, 2024). The distances of potentially multiple disease associations per SLC-SLC pair were integrated into a single distance using the BMA method. The eight modality-specific distances for SLC-SLC pairs varied considerably in range and distribution. Moreover, due to the chosen metrics, the distances for four modalities were continuously distributed, while distances for the other four modalities were truncated to the interval [0,1] (interaction proteomics, subcellular localization, substrate annotation, disease association).

To avoid any distribution-based bias in downstream integration, we transformed the distance distributions to dissimilarity distributions via an Ordered Quantile normalization using the R package ‘bestNormalize’ (version 1.9.1) (Peterson, 2021). Values at the lowest and highest end of the four truncated distance distributions were excluded from this transformation process. The resulting standard normal distributed dissimilarities were further transformed into a truncated normal distribution using the library ‘truncnorm’ (version 1.0-9) (Mersmann *et al*) with µ=0.5 and σ=0.194, chosen so that a corresponding normal distribution would cover 99% of the data in the interval between 0 and 1. Distances at the lower or upper limit from the four modalities with truncated distance distributions were now re-introduced at 0 or 1, respectively (**EV Fig. 5B**). Resulting dissimilarities were subtracted from 1 to result in modality-specific, but more comparable SLC-SLC similarities. These similarities were then combined to a single similarity per SLC pair by a weighted average, putting the highest weight of 3 on structural similarity and the three main experimental data sets in this study (transcriptomics, metabolomics and interaction proteomics), medium weight of 2 on the substrate annotation, and the lowest weight of 1 on tissue expression, subcellular localization and disease association (**Table 2**, **EV Fig. 5D**).

### Constructing the SLC landscape and graph-based clustering

The integrated, pairwise SLC similarities were subtracted from 1 to result in a dissimilarity matrix which in turn was used as the input distance for the construction of a high-dimensional graph representation via the ‘umap-learn’ library (version 0.5.3) (McInnes *et al*, 2018), with a local neighbourhood size of 15. The graph was then embedded into a two-dimensional landscape using 500 training epochs and a minimum distance between embedded points of 0.1.

To identify communities in the SLC landscape, clustering was performed on the high-dimensional graph using the Leiden method, with modularity partition, a resolution of 1.0 and 100 iterations, employing the ‘leiden’ (version 0.4.3.1) and ‘leidenalg’ (version 0.10.1) libraries (Traag *et al*, 2019).

We derived SLC-SLC distances from the high-dimensional graph by defining the distance for each of its edges as its logarithmized adjacency value subtracted from 1 and adding up those edge distances along the shortest path connecting any pair of SLCs. Compared to the weighted-average pairwise SLC-SLC similarity, which was used as the input to the landscape construction, this graph-based distance incorporates local communities as well as global structures in the landscape.

### Analysis of SLC properties in the landscape clusters

To identify drivers of clustering we calculated the adjusted mutual information scores between SLC landscape clusters and a set of discrete SLC properties using ‘sklearn’ library (version 1.3.2) in Python (version 3.11.6) (**EV Fig. 5F**). To find enriched SLC property levels in the clusters, we used the ‘scipy’ library (version 1.13.0) to perform one-sided Fisher’s exact tests for each SLC property levels in each cluster and subsequent multiple-testing correction following the Benjamini-Hochberg procedure (Benjamini & Hochberg, 1995) (**EV Fig. 5G**).

## Acknowledgements

This study received funding from the RESOLUTE and REsolution consortia. RESOLUTE has received funding from the Innovative Medicines Initiative 2 Joint Undertaking under grant agreement No 777372. This Joint Undertaking receives support from the European Union’s Horizon 2020 research and innovation programme and EFPIA. REsolution has received funding from the Innovative Medicines Initiative 2 Joint Undertaking under grant agreement No 101034439. This Joint Undertaking receives support from the European Union’s Horizon 2020 research and innovation programme and EFPIA. This article reflects only the authors’ views and neither IMI nor the European Union and EFPIA are responsible for any use that may be made of the information contained therein.

B.H. was supported by funds from the Austrian Science Fund FWF (DOI:10.55776/KLI1056 to K. Boztug). G.S-F. was supported by the Austrian Academy of Sciences.

## Conflict of interest

G.S-F. is co-founder and owns shares of Solgate GmbH, an SLC-focused company.

D.H. and A.M. are employees and stockholders of Pfizer.

---

**Figure EV1.**
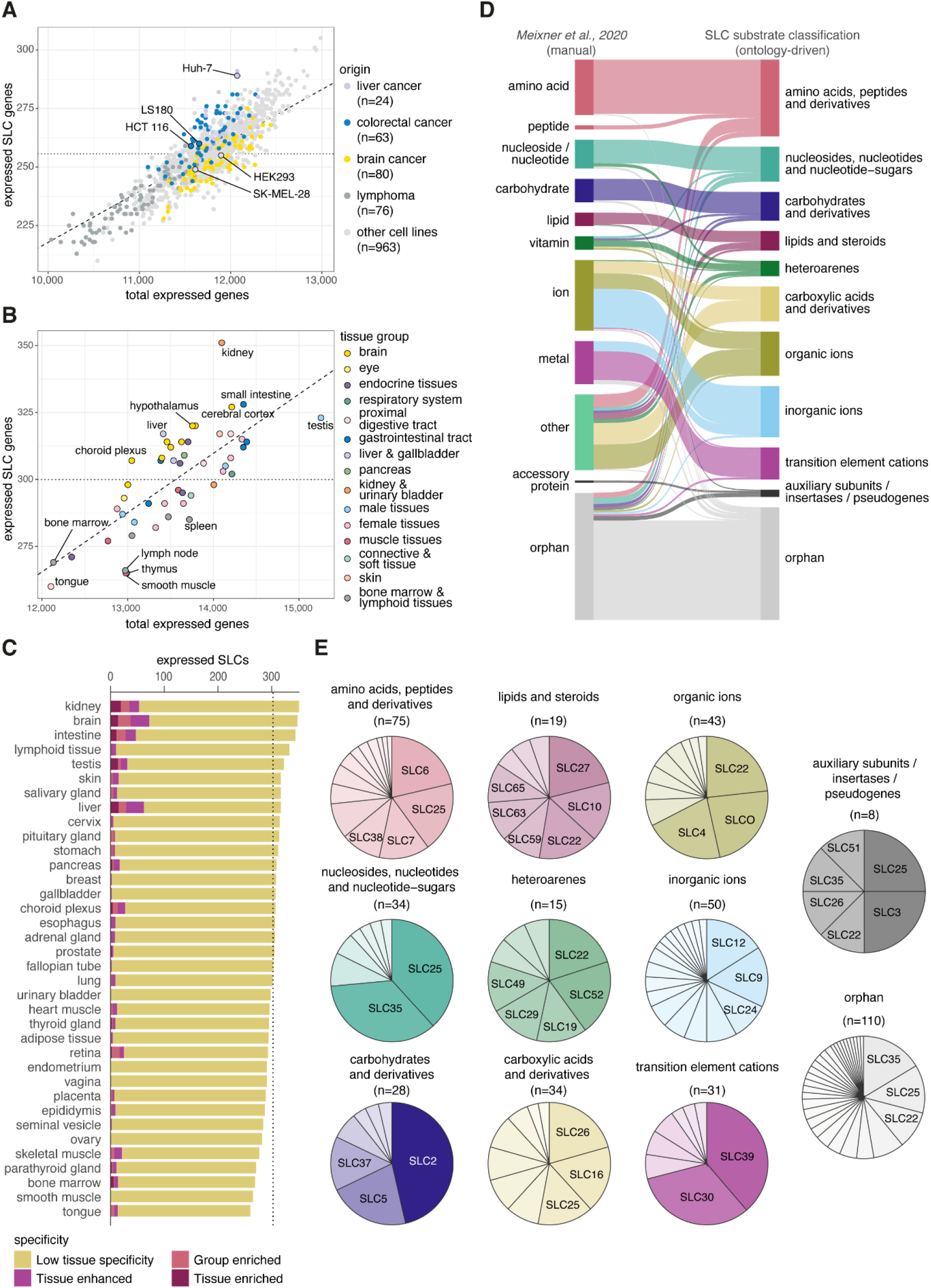
SLC substrate annotation. **(A)** Number of SLC genes expressed in relation to the total number of protein-coding genes expressed in 1,206 human cell lines (data from (Uhlén et al, 2015)). The dotted line indicates the overall average of 256 expressed SLCs per cell line (56.2% of the 455 SLCs with expression data). The dashed line indicates the average of 2.2% of SLC genes among all expressed protein-coding genes in a cell line. Cell lines of selected origin are indicated by colour, and the cell lines employed in the RESOLUTE consortium are labelled. **(B)** Number of SLC genes expressed in relation to the total number of protein-coding genes expressed in 50 human tissues (data from (Uhlén et al, 2015)). The dotted line indicates the overall average of 300 expressed SLCs per tissue (65.9% of the 455 SLCs with expression data). The dashed line indicates the average of 2.2% of SLC genes among all expressed protein-coding genes in a tissue. Tissue groups, as defined in the original data, are indicated by colour and selected tissues deviating from the overall distribution are labelled. **(C)** Total number of SLC genes expressed in 36 tissue groups. The average is indicated by the dotted line. Classification of tissue specificity of gene expression is indicated by colour (data from (Uhlén et al, 2015)). **(D)** Comparison of previous (Meixner et al, 2020) and updated SLC substrate classifications with regards to the terminology and membership of SLCs. The width of the connecting lines corresponds to the number of co-classified SLCs. **(E)** The proportion of SLC families for each substrate class, i.e. number of respective family members compared to the total number of SLCs in each substrate class. The total numbers are given in parentheses.

**Figure EV2.**
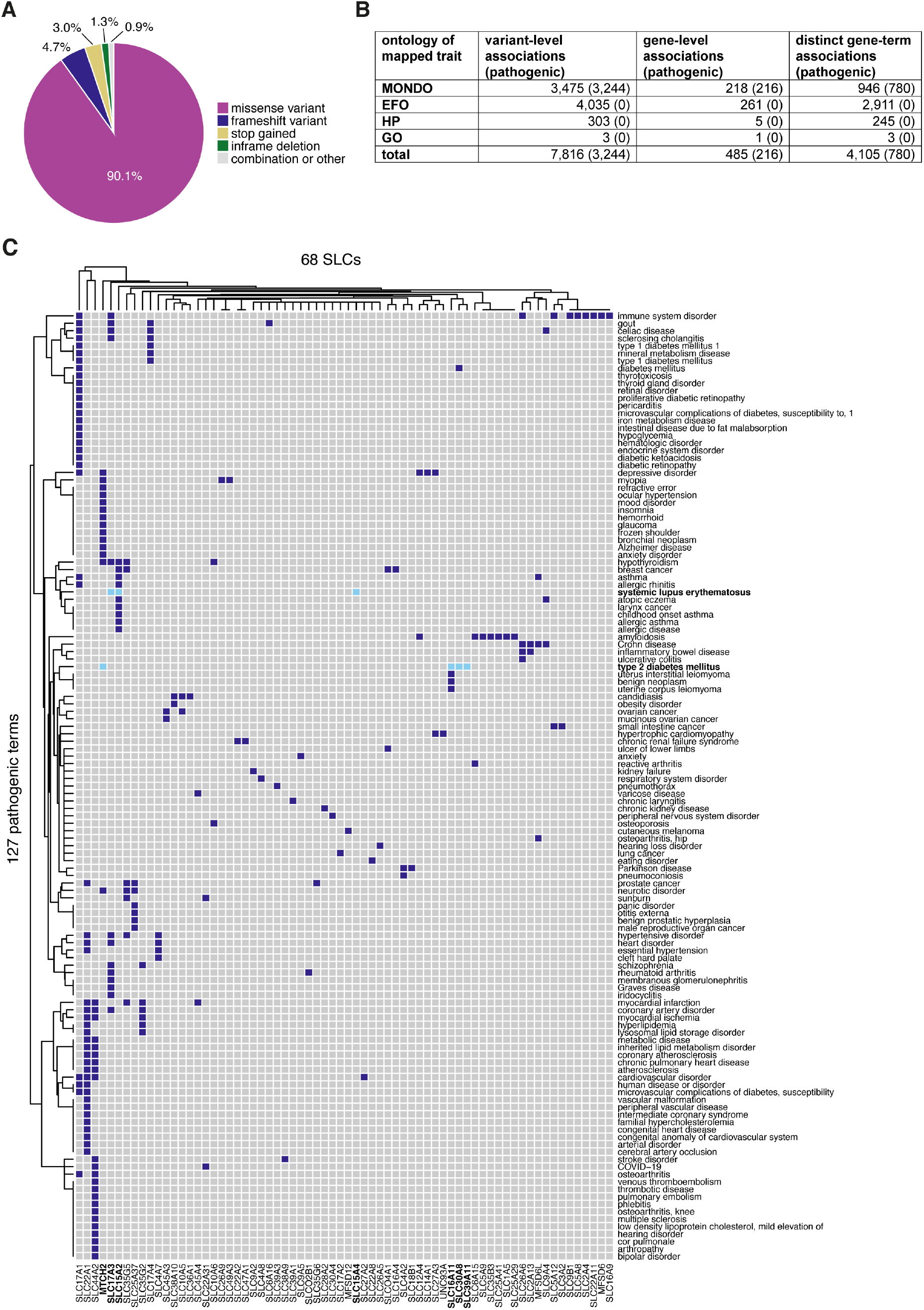
Collection of SLC genetic variants and disease associations. **(A)** Proportions of the main types of genetic alterations of variants with a protein-altering consequence on the canonical transcript of an SLC gene. **(B)** Overview of the number of variants with trait mapping to ontologies Mondo, EFO, HP and GO. Numbers in parentheses are pathogenic variants. **(C)** Heatmap of 68 SLCs without pathogenic associations in ClinVar and Orphanet (x-axis) and novel 127 putative pathogenic terms (y-axis). Examples discussed are marked in bold and light blue.

**Figure EV3.**
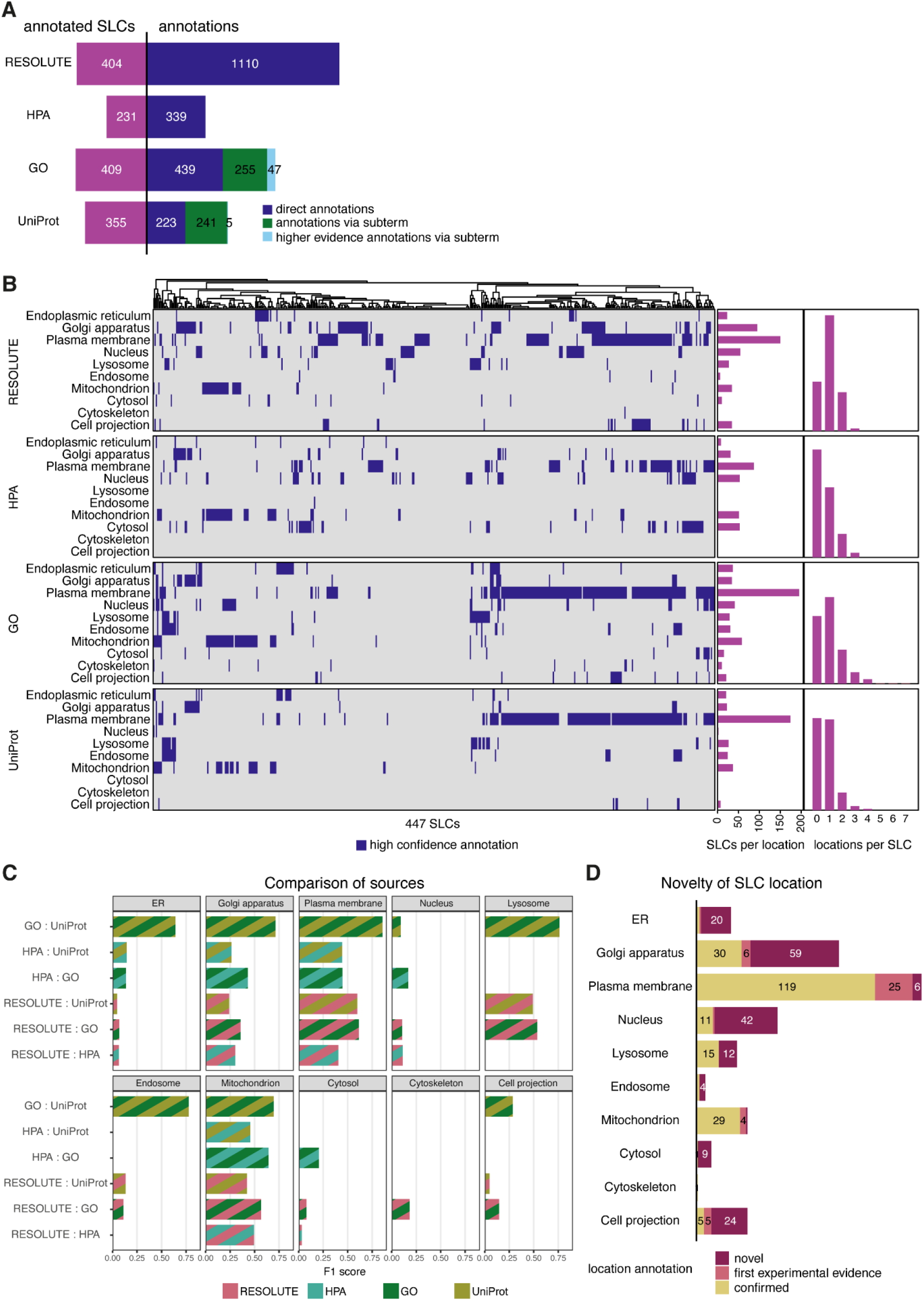
(A) Comparison of the number of location annotations by resource (as in Fig. 3C), highlighting the effect of the described ontology matching process (see Methods). For GO and UniProt, a considerable amount of annotations was only captured via subterms of the 10 selected locations. A smaller number of location annotations did match a selected location term but ended up with an increased evidence level via an additionally mapped subterm of the respective location. **(B)** Overview and comparison of all high-confidence location annotations across RESOLUTE, HPA, GO and UniProt. The histograms shows the prevalence of annotations for each of the selected 10 locations per data source, as well as the distribution of the number of location annotations per SLC for each data source. **(C)** Scoring of the agreement in annotations per subcellular location, for each pair of annotation sources. The F1-score combines precision and recall between the two annotation sources, with a higher score corresponding to higher consistency. **(D)** Novelty of high-confidence experimental localization annotations by the RESOLUTE data set, for each of the 10 selected subcellular locations.

**Figure EV4.**
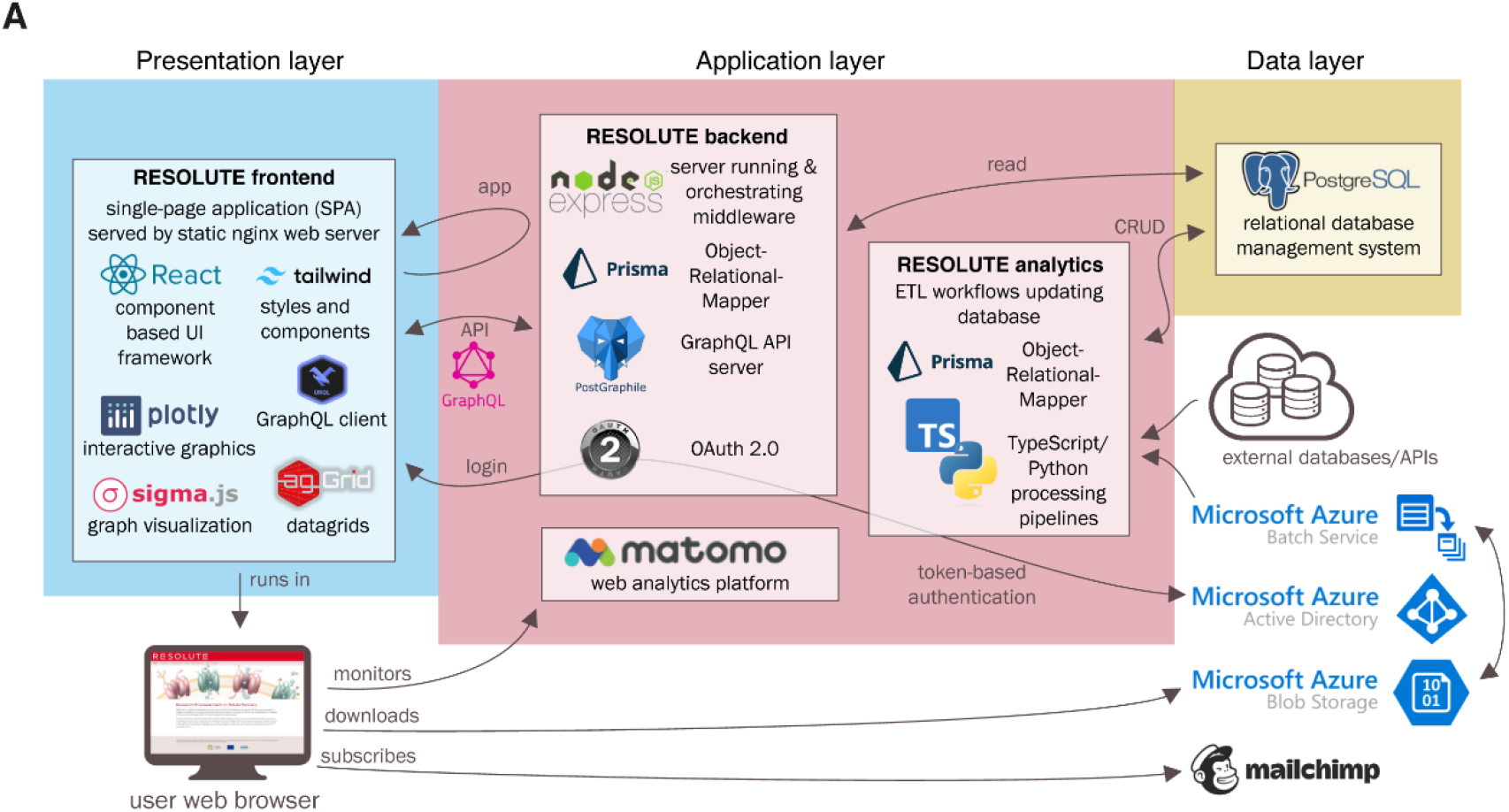
The RESOLUTE web portal software architecture. The web portal features a relational database handling the data layer, a backend and processing workflows handling the application layer, and a fronted in form of a ‘single-page’ web application handling the presentation layer. Please refer to the Methods section for more details. UI: user interface. API: application programming interface. CRUD: create, read, update, delete.

**Figure EV5.**
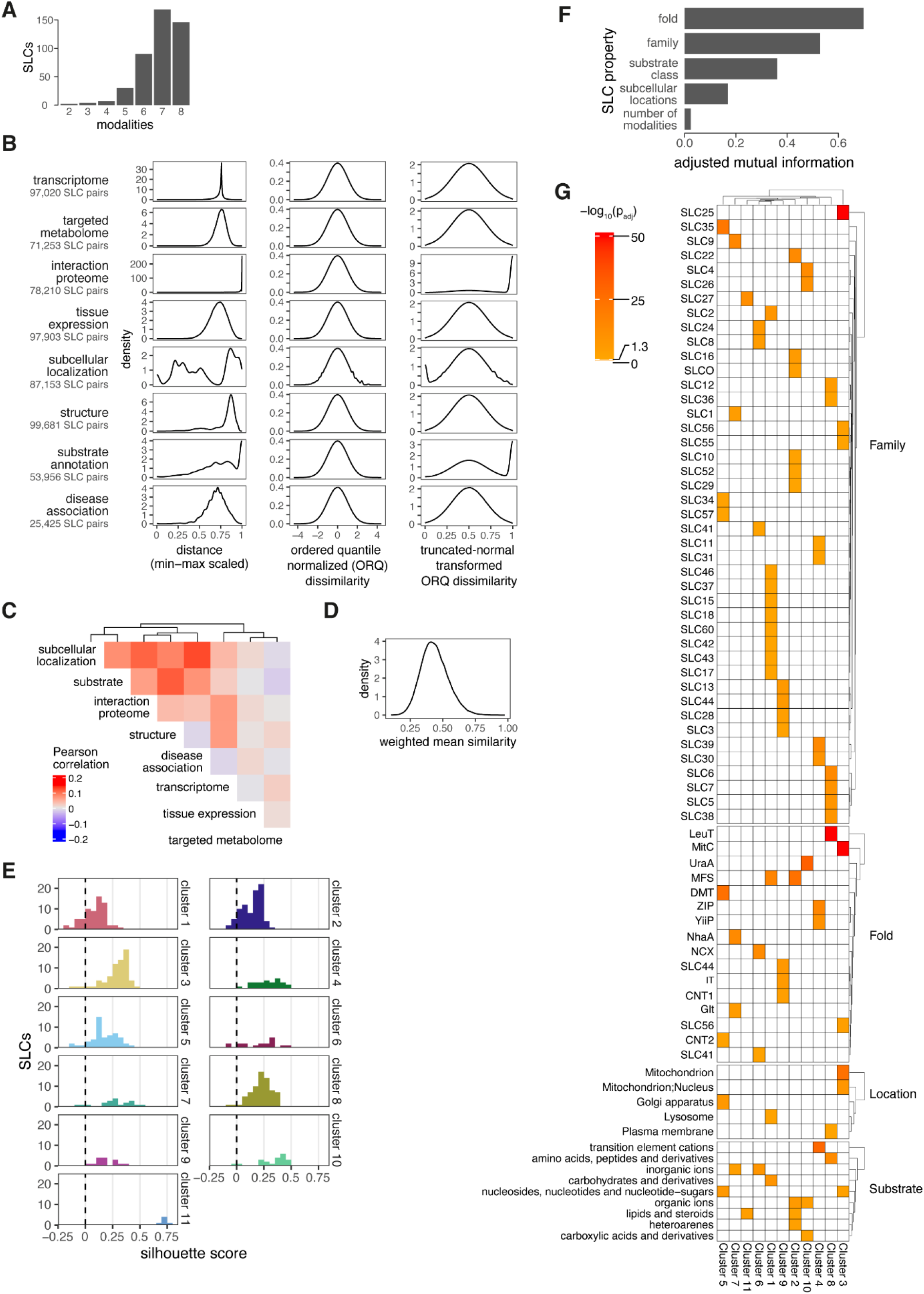
Data sets used in constructing the SLC landscape and analysis of its clustering. **(A)** Distribution of the number of modalities (data sets) available per SLC. **(B)** Distribution of SLC-SLC pair distances per modality. The original distributions of modality-specific distances are shown in the left column, scaled to the interval of [0,1] for visualization purposes. The central column shows the same distributions after transformation to a standard normal distribution using ordered quantile normalization. For distances with an upper or lower limit, values at these limits were excluded from this transformation. The right column shows the normalized distributions transformed to a truncated normal distribution, with the previously excluded values re-introduced at the corresponding boundary, resulting in dissimilarities comparable across modalities. **(C)** Correlations between the eight dissimilarities of all possible SLC-SLC pairs (n=99,681), using Pearson’s method on pairwise- complete observations. **(D)** Distribution of the overall SLC-SLC pair similarities, which resulted from subtracting the weighted average of up to eight dissimilarities for each pair from 1. **(E)** Distribution of silhouette scores of the members of the 11 different clusters derived from the SLC landscape. **(F)** Mutual information analysis of selected, discrete SLC properties. Fold, family, substrate class and localization all share considerable mutual information with the 11 clusters of the SLC landscape, with fold and the fold-related family properties showing the highest correspondence to cluster membership. **(G)** Results of the pairwise enrichment analysis of each SLC property level in each cluster, using Fisher’s exact test. Only SLC property levels with significant enrichments are shown, coloured by their negative log_10_-transformed, multiple-testing corrected p-values.

**Supplemental Table 1.** Listing all 464 human SLC genes with selected properties and annotations. separate Excel file

**Supplemental Table 2.**
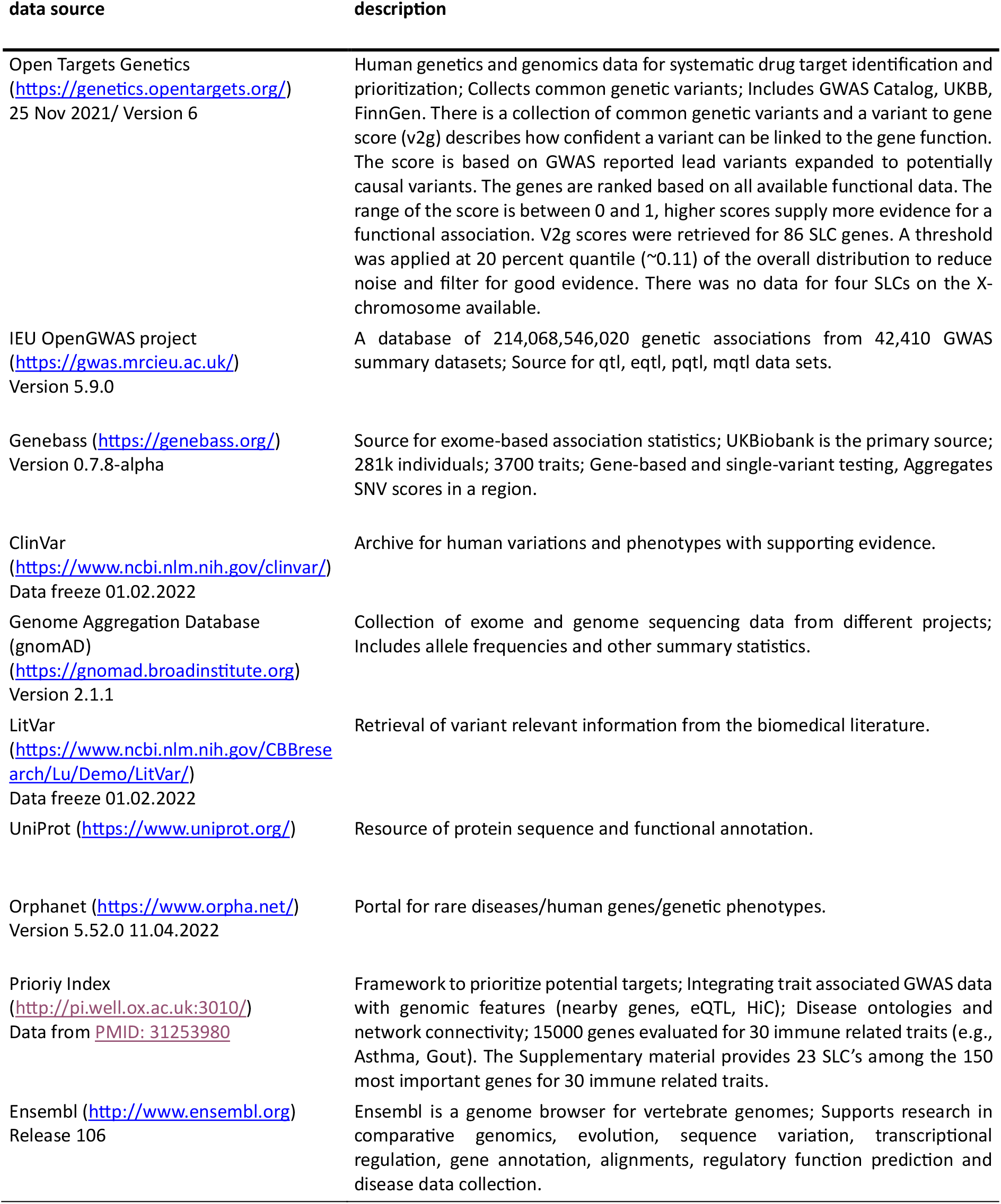
Description of data sources used in survey of SLC superfamily-wide disease associations.

